# Distinct defense strategies against interbacterial antagonism underpin *Acinetobacter* persistence in polymicrobial environments

**DOI:** 10.64898/2026.05.26.727867

**Authors:** Jing Jie, Xin Deng, Meng Zhang, Qingtian Guan, Sanwei Gu, Xia Zhang, Dan Li, Zhao-Qing Luo, Lei Song

**Affiliations:** Department of Respiratory Medicine, Center for Pathogen Biology and Infectious Diseases, State Key Laboratory for Diagnosis and Treatment of Severe Zoonotic Infectious Diseases, The First Hospital of Jilin University; Changchun 130021, China; Bioinformatics Laboratory, Center for Infectious Diseases and Pathogen Biology, The First Hospital of Jilin University, Changchun, China; Cadre’s Ward, The First Hospital of Jilin University, Changchun, China

**Keywords:** Type VI secretion system, polysaccharide capsule, immunity protein, plasmid conjugation, polymicrobial infection

## Abstract

Despite intense interbacterial antagonism mediated by mechanisms such as the type VI secretion system (T6SS), pathogenic *Acinetobacter* frequently persist within highly competitive polymicrobial infections. How these pathogens navigate such hostile environments to achieve co-existence remains poorly understood. Here, we show that pathogenic *Acinetobacter* employ distinct, multifaceted strategies to resist elimination by both distantly related competitors and closely related kin. T6SS-active strains outcompete heterologous bacteria by deploying a large and diverse repertoire of antibacterial effectors. In contrast, some naturally T6SS-deficient strains resist exogenous T6SS attacks by elaborating a unique, high-density polysaccharide capsule (KL34) that functions as an effective physical shield. Most intriguingly, we uncover a mechanism that prevents lethal competition among kins. A conjugative multidrug-resistant plasmid encoding T6SS repressors is transferred into aggressive strains upon T6SS-mediated attack, leading to suppression of their antibacterial activity. We also found that conjugation is induced by T6SS attack, which effectively suppresses attacking kin and promotes co-infection *in vivo*. Together, these findings reveal a dynamic and multilayer defense program that enables *Acinetobacter* to withstand interbacterial warfare while facilitating co-existence. Our study establishes a paradigm in which damage-triggered plasmid transfer enforces targeted suppression, reshaping microbial interactions in polymicrobial communities.

## Introduction

*Acinetobacter* species, most notably *Acinetobacter baumannii,* are common opportunistic pathogens responsible for severe hospital-acquired infections, including pneumonia, bloodstream infections, and wound sepsis. Notorious for their multidrug resistance and environmental resilience, these bacteria are incredibly difficult to eradicate in clinical settings ^1^. A defining, yet complicating, feature of *Acinetobacter* infections is their highly polymicrobial nature. *Acinetobacter* is frequently co-isolated alongside other nosocomial pathogens, such as *Klebsiella pneumoniae* or *Pseudomonas aeruginosa*. Such mixed-species infections are often more severe and recalcitrant to treatment than mono-infections ^2-4^. Furthermore, multiple distinct *Acinetobacter* strains or closely related species such as *A. nosocomialis* and *A. pittii*, are frequently documented to simultaneously colonize a single patient, forming a complex intra-genus consortium *in vivo* ^5,6^. These clinical observations raise a fundamental ecological question: how do pathogenic *Acinetobacter* survive and thrive when surrounded by a gamut of lethal microbial competitors, including their own kin?

Within these dense polymicrobial communities, bacteria are engaged in a constant arms race for niche dominance. Many Gram-negative bacteria deploy the type VI secretion system (T6SS) as a weapon to eliminate neighboring microbes ^7-9^. The T6SS is a spike-like contractile nanomachine that injects toxic effectors directly into rival bacteria in a contact-dependent manner ^10^. In pathogen-rich environments, such T6SS-mediated warfare dictates community composition. *A. baumannii* and its relatives possess a functional T6SS that is primarily dedicated to interbacterial competition rather than host infection ^9,11,12^.

However, pathogenic *Acinetobacter* presents a fascinating paradox regarding interbacterial defense. While many clinical isolates can employ the T6SS defensively and offensively to kill competitors ^9,11,13^, a significant fraction of *Acinetobacter* strains naturally lack an intact T6SS locus or fail to express a functional system ^14-16^. Moreover, some strains harbor mobile genetic elements, such as the multidrug-resistant plasmid pAB3, which actively repress T6SS expression ^17,18^, implying that these bacteria actively regulate the activity of their weapons during infection. This paradox creates two pressing mechanistic questions: First, how do T6SS-deficient Acinetobacter strains survive assaults from T6SS-active competitors? Second, when multiple *Acinetobacter* species co-colonize a host, how do they prevent T6SS-mediated fratricidal effects to achieve co-existence?

In this study, we demonstrate that pathogenic *Acinetobacter* navigate polymicrobial conflicts by deploying a multi-tiered, highly dynamic defense strategy. Using clinical isolates and prototypical Gram-negative pathogens as models, we found that while some strains outgun their distant competitors using an unusually expansive T6SS effector repertoire, naturally T6SS-deficient strains survive by elaborating a unique, high-density polysaccharide capsule (KL34) that functions as an effective physical shield. Moreover, intriguingly, these bacteria actively share a T6SS-repressing plasmid to avoid killing among kins and conjugative dissemination of this plasmid is induced by T6SS-mediated attack. These events effectively suppress the aggressors and promote cooperative polymicrobial infection *in vivo*. Together, our findings shed light on the sophisticated and underappreciated mechanisms that enable *Acinetobacter* to persist and cohabit in intensely competitive clinical environments.

## Results

### Protection against Gram-negative competitors via abundant T6SS effector repertoire

We determined the profile of microbial communities in 873 blood, sputa or bronchoalveolar lavage fluid (BALF) samples collected from patients with pulmonary infections by next-generation sequencing (NGS), and found that pathogenic *Acinetobacter* spp., including *A. baumannii*, *A. nosocomialis*, and *A. pittii* were present frequently in polymicrobial infections. Specifically, infections caused solely by *Acinetobacter* comprised only 21.99% of the samples. In contrast, a striking 78.01% of the cases were polymicrobial infections (**Figure S1A**). Among these polymicrobial cases, the most frequently co-occurring bacterial pathogens with *Acinetobacter* spp. were *K. pneumoniae* (41.43%), *Stenotrophomonas maltophilia* (34.41%), and *Enterococcus faecalis* (28.70%) (**Figure S1B**). These findings reveal that pathogenic *Acinetobacter* spp. frequently coexist with other bacterial pathogens in patients, which raises a clinically relevant question: how do the bacterium survive from T6SS-mediated attacks from co-infecting bacteria?

To investigate whether pathogenic *Acinetobacter* employ T6SS to protect themselves during polymicrobial infections, we selected clinical isolates of *K. pneumoniae*, *Enterobacter cloacae*, and *Serratia marcescens*, each with active T6SS evidenced by robust Hcp secretion (**Figure 1A**) and effective killing of *Escherichia coli* for analysis (**Figure 1B**). To assess the defense capacity of pathogenic *Acinetobacter*, we used three clinical isolates from polymicrobial infections — *A. baumannii* Ab12, *A. nosocomialis* An25, and *A. pittii* Ap42, each with an active T6SS and their respective Δ*tssM* mutants defective in T6SS function, as preys (**Figure 1C**). Results from pair-wise competition experiments showed that wild-type Ab12, An25, and Ap42 but not their Δ*tssM* mutants were resistant to killing by the Gram-negative predators (**Figure 1D-E**). Conversely, when pathogenic *Acinetobacter* isolates were tested as predators, they displayed robust T6SS-mediated killing of *K. pneumoniae*, *E. cloacae*, and *S. marcescens* (**Figure 1F-G**). These results suggest that T6SS allows pathogenic *Acinetobacter* isolates to attack, thus effectively defending against competing bacteria.

**Figure 1.**
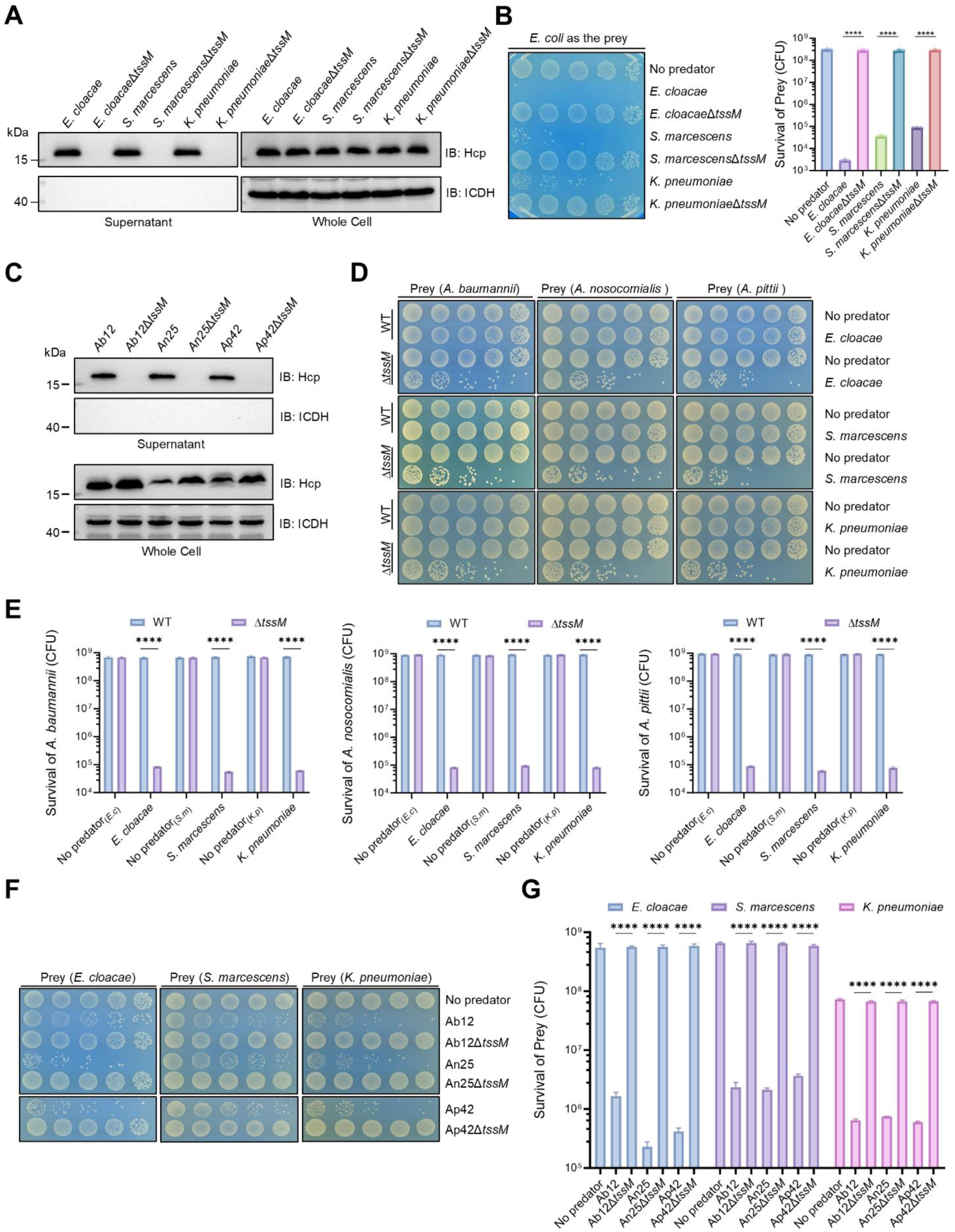
The T6SS mediates both offense and defense in interbacterial competition among pathogenic *Acinetobacter* strains. **(A)** Hcp secretion by clinical isolates of *K. pneumoniae*, *E. cloacae*, and *S. marcescens*. The presence of Hcp in the culture supernatant indicates an active T6SS. The cytosolic protein isocitrate dehydrogenase (ICDH) was probed as a loading control. **(B)** Killing of *E. coli* by Gram-negative pathogens. *E. coli* cells were mixed with the indicated testing strains at a 1:1 ratio and co-incubated on LB agar for 4 h. The survival of *E. coli* was assessed by plating serial dilutions on selective media. Representative images of surviving colonies (left) and the viable counts of surviving prey cells (right) are shown. **(C)** T6SS activity in *Acinetobacter* clinical isolates determined by Hcp secretion. Proteins from the culture supernatants of wild-type *A. baumannii* Ab12, *A. nosocomialis* An25, *A. pittii* Ap42, and their corresponding T6SS-defective mutants (Δ*tssM*) were separated by SDS-PAGE and probed for Hcp via immunoblotting. **(D-E)** Survival of *Acinetobacter* strains during competition with distantly related Gram-negative predators. *E. cloacae*, *S. marcescens*, or *K. pneumoniae* were mixed with the indicated wild-type *Acinetobacter* strains or their Δ*tssM* mutants as described in (B). Representative images (D) and quantitative survival results (E) are shown. **(F-G)** Survival of *E. cloacae*, *S. marcescens*, and *K. pneumoniae* during competition with *Acinetobacter* strains. Experiments were performed as described in (D). Representative images (F) and quantitative survival results (G) are shown. For (B), (E), and (G), quantitative data are presented as the mean ± SEM of three independent biological replicates. Statistical significance was determined using an unpaired, two-tailed Student’s *t*-test on log_10_-transformed CFU values (****, *P* < 0.0001).

To identify the T6SS-related factors that allow pathogenic *Acinetobacter* spp. to outcompete other clinically important pathogens such as *K. pneumoniae*, *E. cloacae*, and *S. marcescens*, we hypothesized that the breadth of their effector arsenal may provide a competitive advantage. Comparative genomic analysis using NCBI genome datasets revealed that the average number of VgrG proteins encoded by pathogenic *Acinetobacter* species is significantly higher than those of *Klebsiella*, *Enterobacter*, *Neisseria*, or *Serratia* species, with counts only slightly lower than those found in *Pseudomonas* spp. (**Figure 2A**, top and bottom panels). Given that VgrG abundance often correlates with the diversity of the effector repertoire ^19^, these results suggest that pathogenic *Acinetobacter* spp., exemplified by *A. baumannii*, is equipped with more T6SS effectors than other often co-residing Gram-negative pathogens.

**Figure 2.**
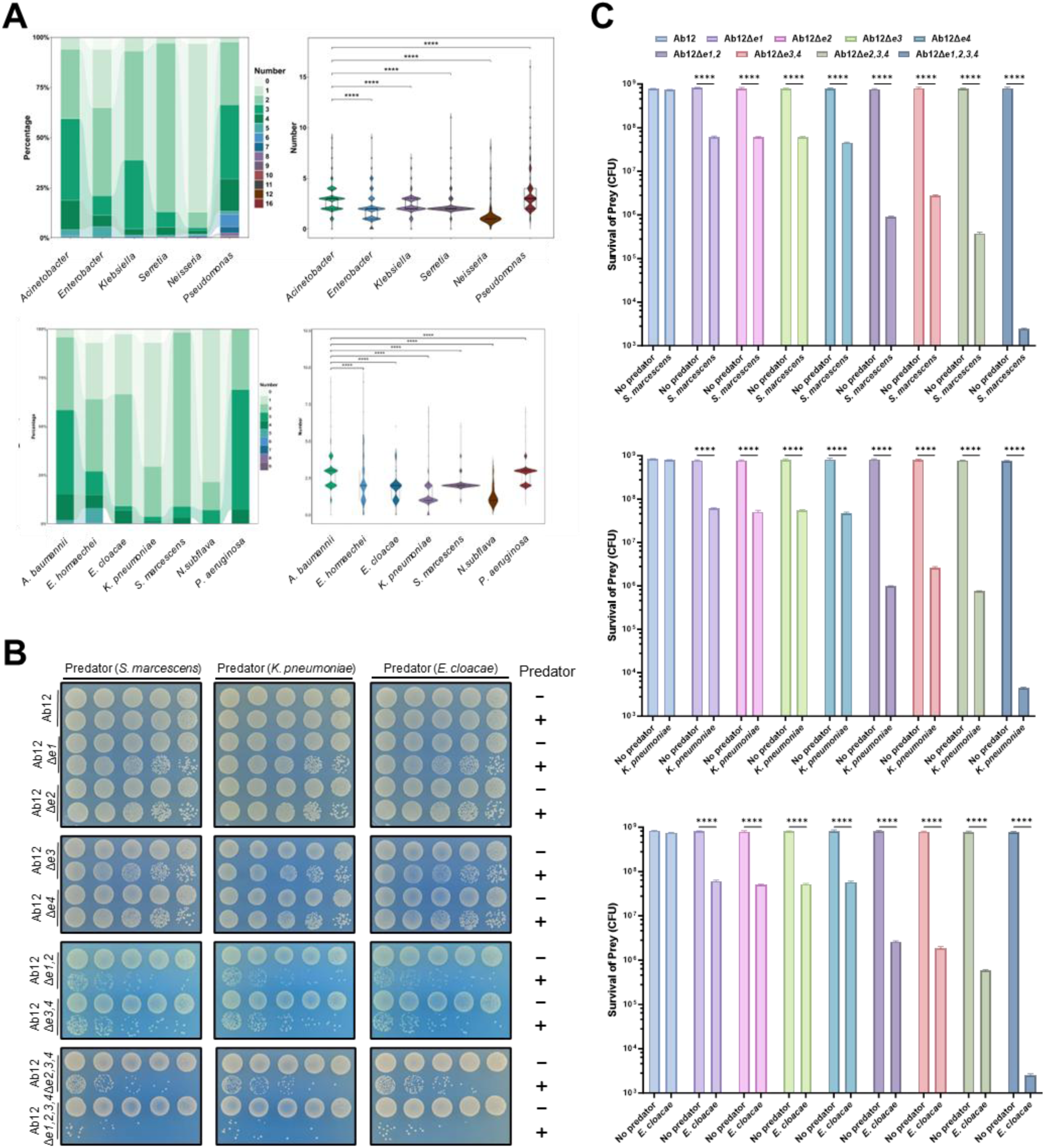
An expansive effector repertoire is required for pathogenic *Acinetobacter* to outcompete Gram-negative pathogens. **(A)** Comparative genomic analysis of T6SS arsenals across multiple bacterial taxa. The distribution of VgrG proteins was analyzed at the genus level (top panel) and across selected clinically important species (bottom panel). Violin plots demonstrate that pathogenic *Acinetobacter* encode a significantly higher average number of VgrG proteins than most other analyzed groups, with counts only slightly lower than those found in *Pseudomonas* spp. **(B-C)** An intact effector repertoire is essential for *A. baumannii* to defend against T6SS-mediated killing. Wild-type *A. baumannii* Ab12 and its derivatives lacking single or combinatorial effector genes (ranging from single Δ*e* mutants to the quadruple mutant Δ*e1,2,3,4*) were subjected to competition against *S. marcescens*, *K. pneumoniae*, or *E. cloacae*. Representative images of surviving *A. baumannii* (B) and quantitative survival results (C) are shown. Note that the Δ*e1,2,3,4* mutant exhibited the highest susceptibility to competitor attacks. For (C), quantitative data are presented as the mean ± SEM of three independent biological replicates. Statistical significance was determined using a one-way ANOVA followed by Dunnett’s multiple comparisons test against the wild-type Ab12 control on log_10_-transformed CFU values (****, *P* < 0.0001).

To validate the contribution of these effectors in inter-species competition, we examined clinical isolate AB12, which harbors four VgrG-effector pairs (**Figure S2**). We constructed a panel of mutant strains lacking various combinations of effector genes (e.g., Δ*e1*; Δ*e1,2*; Δ*e1,3*) as well as a strain lacking all four effector genes (Δ*e1,2,3,4*). In interbacterial competition assays where *A. baumannii* AB12 or its mutants were mixed with strains of *K. pneumoniae*, *E. cloacae* or *S. marcescens*, the wild-type *A. baumannii* strain was highly resistant to killing, while these effector-deficient mutants displayed susceptibility to competitor attack, with the degree of killing correlating with the number of effectors deleted. Among these, the Δ*e1,2,3,4* strain exhibited the greatest sensitivity (**Figure 2B-C**), underscoring the cumulative effects of these effectors in resisting T6SS-mediated assaults from competing bacteria. These results suggest that the expansive repertoire of VgrG-associated effectors endows the T6SS of pathogenic *Acinetobacter* spp. to outcompete other Gram-negative pathogens.

### Resistance to T6SS attack by a unique capsule

Although pathogenic *Acinetobacter* spp. generally relies on potent T6SS activity to defend against Gram-negative competitors, genomic surveys revealed a paradox: at least 16.2% of *A. baumannii* genomes in public databases do not encode a complete T6SS cluster. This ratio is markedly higher than those observed in *P. aeruginosa, E. cloacae,* or *S. marcescens*, and is only slightly lower than that of *K. pneumoniae* and *N. subflava* (**Figure S3**). Consistent with this, among the 186 strains of our clinical *Acinetobacter* isolates tested, only 67 strains (36%) secreted Hcp and exhibited T6SS-dependent killing of *E. coli* (**Figure S4A-C)**.

We next randomly tested 44 T6SS-inactive clinical isolates (Hcp-negative, unable to kill *E. coli*; some representatives were shown in **Figure 3A**) for the ability to compete against T6SS-competent strains of *K. pneumoniae, E. cloacae* or *S. marcescens*. While the majority of these strains (38 out of 44) were highly sensitive to killing by these bacteria, several strains exhibited strong resistance (**Figure 3B-C**). These findings suggest that some pathogenic *Acinetobacter* strains exhibit resistance to T6SS attack by mechanisms not related to T6SS activity.

**Figure 3.**
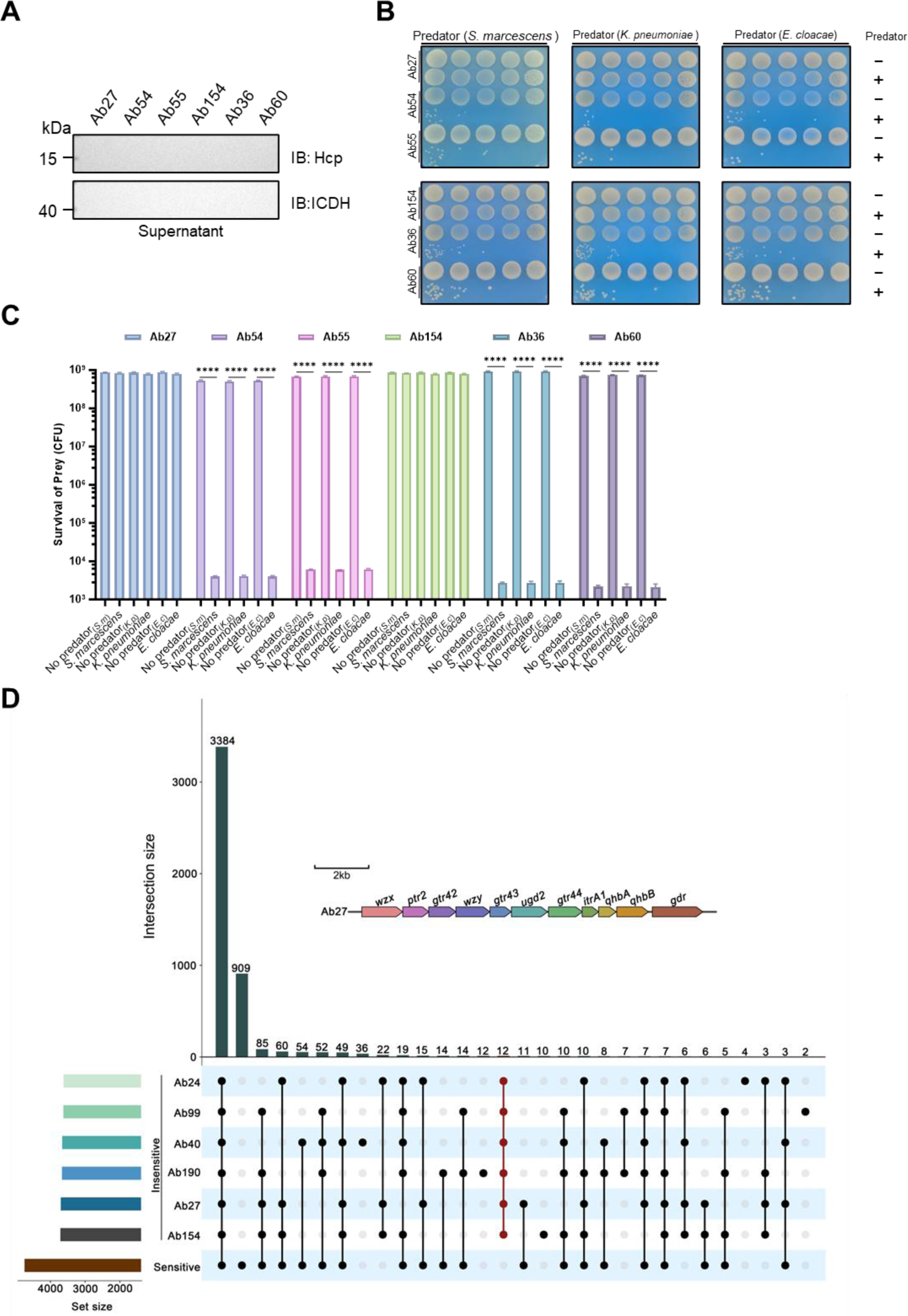
A specific capsule biosynthesis gene cluster governs the resistance of T6SS-inactive *Acinetobacter* isolates against exogenous attacks. **(A)** Hcp secretion in representative clinical *A. baumannii* isolates. Strains Ab27, Ab54, Ab55, Ab154, Ab36, and Ab60 lacked detectable Hcp secretion, indicating a naturally inactive T6SS. ICDH served as a loading control. **(B-C)** Susceptibility of T6SS-inactive isolates to Gram-negative competitors. A total of 44 Hcp-negative isolates were tested as prey against T6SS-active *S. marcescens*, *K. pneumoniae*, or *E. cloacae*. Representative images (B) and quantitative results (C) reveal two distinct phenotypic groups: susceptible strains (e.g., Ab54, Ab55, Ab36, Ab60; n=38) and strongly resistant strains (e.g., Ab27, Ab154; n=6). For (C), quantitative data are presented as the mean ± SEM of three independent experiments performed in triplicate. Statistical significance comparing predator-exposed groups to the respective "No predator" controls was determined using an unpaired, two-tailed Student’s *t*-test on log_10_-transformed CFU values (****, *P* < 0.0001). **(D)** Comparative genomic identification of a resistance-associated genetic signature. The UpSet plot visualizes the intersection of gene sets across resistant (Insensitive) and sensitive strains. A specific 12-gene cluster uniquely shared among the resistant isolates is highlighted (red line). The inset illustrates the genetic architecture of this cluster in strain Ab27, which encodes the KL34-type capsule biosynthesis pathway (e.g., *wzx, gtr42, wzy, gtr43*).

Comparative genomic analysis revealed that these resistant strains harbor a cluster of 12 genes that may be responsible for this phenotype (**Figure 3D**). Notably, 11 of these genes predicted to encode biosynthesis proteins for type KL34 capsule are organized into an operon-like structure. Capsule typing and phylogenetic analysis of our entire collection revealed that all *Acinetobacter* strains resistant to T6SS killing possess genes for the production of the KL34 capsule (**Figure S5**). Conversely, T6SS-deficient *Acinetobacter* strains (lacking Hcp secretion, **Figure S6A**) harboring genes for other capsule types were all susceptible to killing by *K. pneumoniae*, *E. cloacae* or *S. marcescens* at different levels (**Figure S6B-C**). These results demonstrate that a subset of T6SS-defective strains has evolved to elaborate a unique capsule to confer resistance to T6SS assaults from other Gram-negative bacteria.

We further analyzed the resistance of strain Ab27 (KL34 type) against *E. cloacae* by mixing it with wild-type or its Δ*tssM* mutant and performed transcriptomic analysis. Our results revealed changes in the expression of a large number of genes in Ab27 upon exposure to wild-type *E. cloacae*. Importantly, multiple genes within the KL34 capsule biosynthesis locus, including *gtr42, ugd2, ptr2, gtr44, gtr43,* and *wzy*, were significantly upregulated (**Figure 4A**). Such T6SS-dependent induction was also observed when the changes were determined by qPCR (**Figure 4B**).

**Figure 4.**
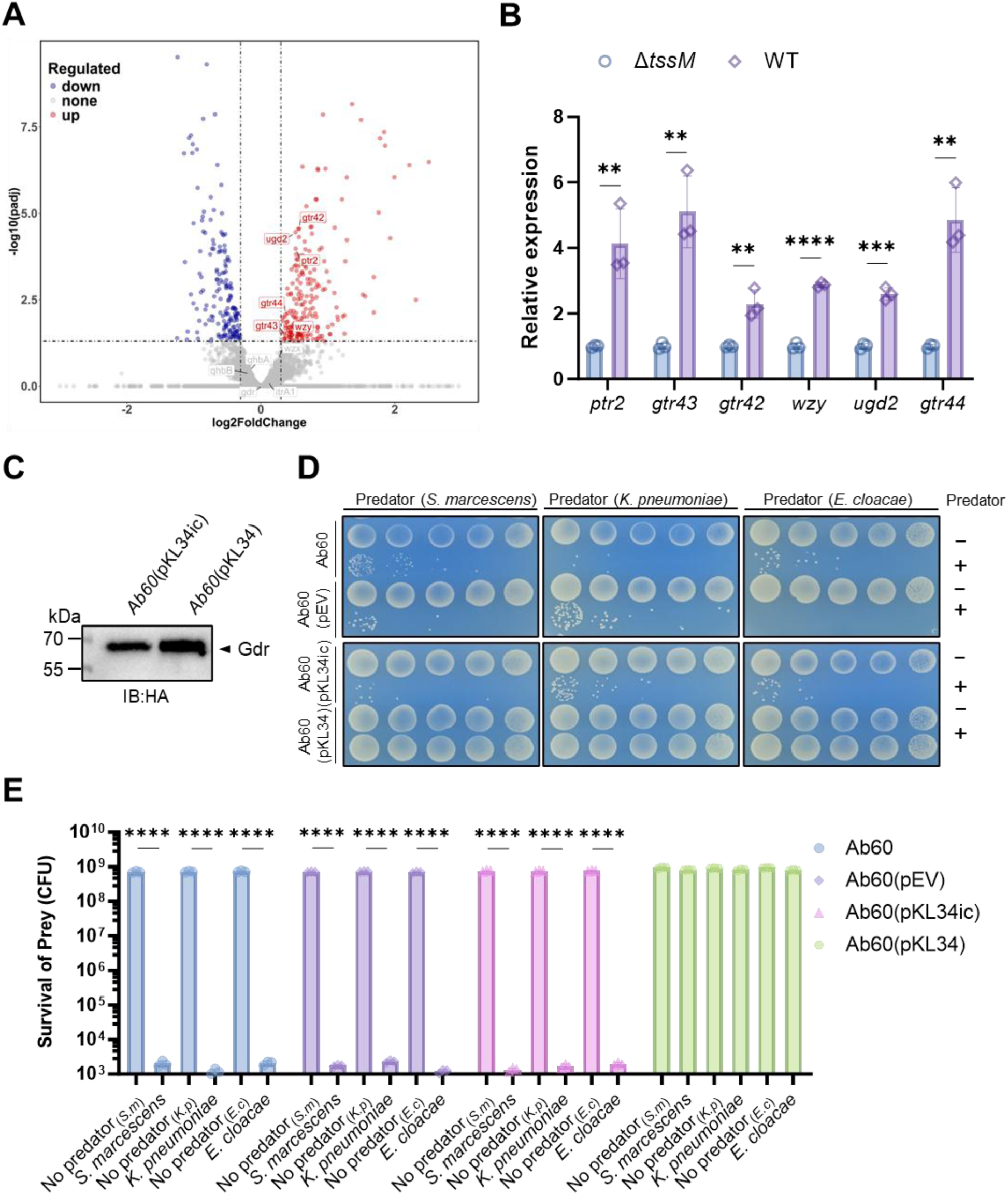
Transcriptional activation of the KL34 capsule cluster upon T6SS attack is essential for physical defense. **(A)** Transcriptomic reprogramming of the resistant strain *A. baumannii* Ab27. Gene expression profiles were compared between Ab27 co-incubated with wild-type *E. cloacae* (T6SS-active) versus its T6SS-inactive Δ*tssM* mutant. The volcano plot highlights significantly upregulated genes (red dots), encompassing multiple genes within the KL34 capsule biosynthesis locus (e.g., *gtr42, ugd2, ptr2, gtr44, gtr43*, and *wzy*). **(B)** RT-qPCR validation of T6SS-dependent transcriptional induction in Ab27. Relative mRNA levels were measured following co-incubation with wild-type *E. cloacae* compared to the Δ*tssM* mutant. **(C)** Expression validation of the KL34 gene cluster in the sensitive background strain Ab60. Derivatives of Ab60 harboring plasmids with either the complete 11-gene KL34 cluster (pKL34) or a partial cluster lacking the *ptr2* gene (pKL34ic) were constructed. An HA epitope was appended to the distal *gdr* gene to confirm proper expression. **(D-E)** The complete KL34 cluster is sufficient to confer resistance to T6SS-mediated killing. Derivatives of Ab60 carrying the empty vector (pEV), pKL34ic, or pKL34 were used as prey against *S. marcescens*, *K. pneumoniae*, or *E. cloacae*. Ab27 and wild-type Ab60 served as resistant and sensitive controls, respectively. Representative images (D) and quantitative results (E) are shown. Quantitative data in (B) and (E) are presented as the mean ± SEM of three independent biological replicates. Statistical significance in (B) was determined by unpaired, two-tailed Student’s *t*-tests (**, *P* < 0.01; ***, *P* < 0.001; ****, *P* < 0.0001). Statistical significance in (E) was determined by a one-way ANOVA followed by Tukey’s multiple comparisons test on log_10_-transformed CFU values (****, *P* < 0.0001).

Beyond the capsule locus, this analysis revealed a global defensive reprogramming. We identified the significant upregulation of genes involved in the synthesis of poly-beta-1,6-N-acetyl-D-glucosamine (PGA) biofilm matrix (e.g., *pgaB*, *pgaC*, *pgaD*) and extracellular polysaccharides (*epsO*, *epsJ*). Notably, we also observed the upregulation of a suite of T6SS structural and regulatory genes (e.g., *tssB*, *tssC*, *tssF*, *tssM*, and *hcp*), indicative of a damage-sensing "tit-for-tat" counterattack response, despite Ab27 being naturally T6SS-silent under monoculture conditions. Concurrently, the attacked cells exhibited a robust iron-scavenging response—characterized by the upregulation of TonB-dependent receptors and siderophore biosynthesis clusters—alongside the activation of oxidative stress defense mechanisms (**Supplementary Table S1**).

To validate the role of the KL34 gene cluster in resistance to T6SS attack, we constructed two plasmids, one harboring the complete set of the KL34 biogenesis genes and one carrying a DNA fragment lacking the *ptr2* gene. These plasmids were introduced into the T6SS-sensitive *A. baumannii* clinical strain Ab60, to generate Ab60(pKL34) and Ab60(pKL34ic), respectively. To facilitate gene expression detection, an HA tag was appended to the C-terminus of the *gdr* gene distal in the operon. The operon was properly expressed in strains Ab60(pKL34) and Ab60(pKL34ic) (**Figure 4C**), but only strain Ab60(pKL34) exhibited resistance to killing by *K. pneumoniae*, *E. cloacae* or *S. marcescens* (**Figure 4D-E**). These complementation results firmly establish that while *Acinetobacter* mounts a multifaceted global alarm response involving biofilms and T6SS counterattacks, the high-density KL34 capsule serves as the primary and indispensable physical shield to withstand exogenous T6SS assaults.

We also compared the capsule morphology between the T6SS-resistant KL34-type strains (Ab27, Ab40, and Ab154) and the T6SS-sensitive strains (Ab60 [KL2-type], Ab85 [KL84-type], and Ab111 [KL105-type]) using transmission electron microscopy (TEM). This analysis revealed that while the KL34 capsules were significantly thinner than that of the KL2-type strain Ab60, their thickness was comparable to those of the KL84 and KL105 strains (**Figure S7A-B**). Importantly, the capsular layer of the KL34 strains exhibited a more compact capsule that retained more of the staining agent than all other tested T6SS-sensitive strains, regardless of their overall capsule thickness (**Figure S7C**). Furthermore, expression of the KL34 capsule gene set allowed strain Ab60 to elaborate a capsule resembling that of strain Ab27 (**Figure S8A-C**). These results indicate that capsule density, rather than thickness, is the key feature required for protection against T6SS attack.

### Plasmid dissemination universally silences T6SS to prevent intra-genus antagonism

Acinetobacter frequently coexists with closely related species in polymicrobial infections ^5,6^, suggesting that these bacteria have evolved mechanisms to avoid intra-genus killing. We envisioned that such mechanisms are likely involved in regulation of T6SS expression and performed phylogenetic analysis of pathogenic and evolutionarily distinct nonpathogenic *Acinetobacter* species (**Figure S9A**). These analyses revealed that pathogenic species possess a uniquely organized T6SS locus (Type I), which differs in the orientation of the structural genes from loci found in non-pathogenic species (Type II) (**Figure S9B**). Furthermore, the promoter regions of Type I feature highly conserved –35 and –10 elements, and a palindromic motif (**Figure S10A**). This conserved promoter architecture implies a shared regulatory mechanism among pathogenic lineages.

The multidrug-resistant plasmid pAB3 has been shown to repress T6SS in *A. baumannii* via two TetR-family repressors, TetR1 and TetR2 ^17^. Given the conserved promoters, we hypothesized that this repression is a universal phenomenon across pathogenic *Acinetobacte*r species. Indeed, expression of TetR1 or TetR2 not only inhibited Hcp synthesis in diverse *A. baumannii* clinical isolates but also robustly suppressed Hcp expression in a few tested pathogenic species, including *A. nosocomialis*, *A. pittii*, and *A. seifertii* (**Figure S10B**). As expected, these repressors had no impact on the T6SS activity of non-Acinetobacter pathogens (e.g., *E. cloacae* or *K. pneumoniae*) (**Figure S10C**). Importantly, the natural acquisition of the entire pAB3 plasmid via its Dot/Icm-like Type IV secretion system (T4SS) ^20^ completely abolished Hcp expression in these recipient pathogenic species (**Figure S10D**).

The ability of the pAB3 plasmid to universally silence T6SS expression across diverse pathogenic Acinetobacter species prompted us to examine whether it actively confers protection against T6SS attacks from kin strains. Plasmidless *A. nosocomialis* strain An25, *A. pittii* strain Ap42, and *A. seifertii* strain As86 each effectively killed *A. baumannii* strains Ab55 and Ab60, both of which are defective in Hcp secretion (**Figure 5A**). Strikingly, introduction of wild-type pAB3 into Ab55 and Ab60 rendered them highly resistant to such killing (**Figure 5B-C**).

**Figure 5.**
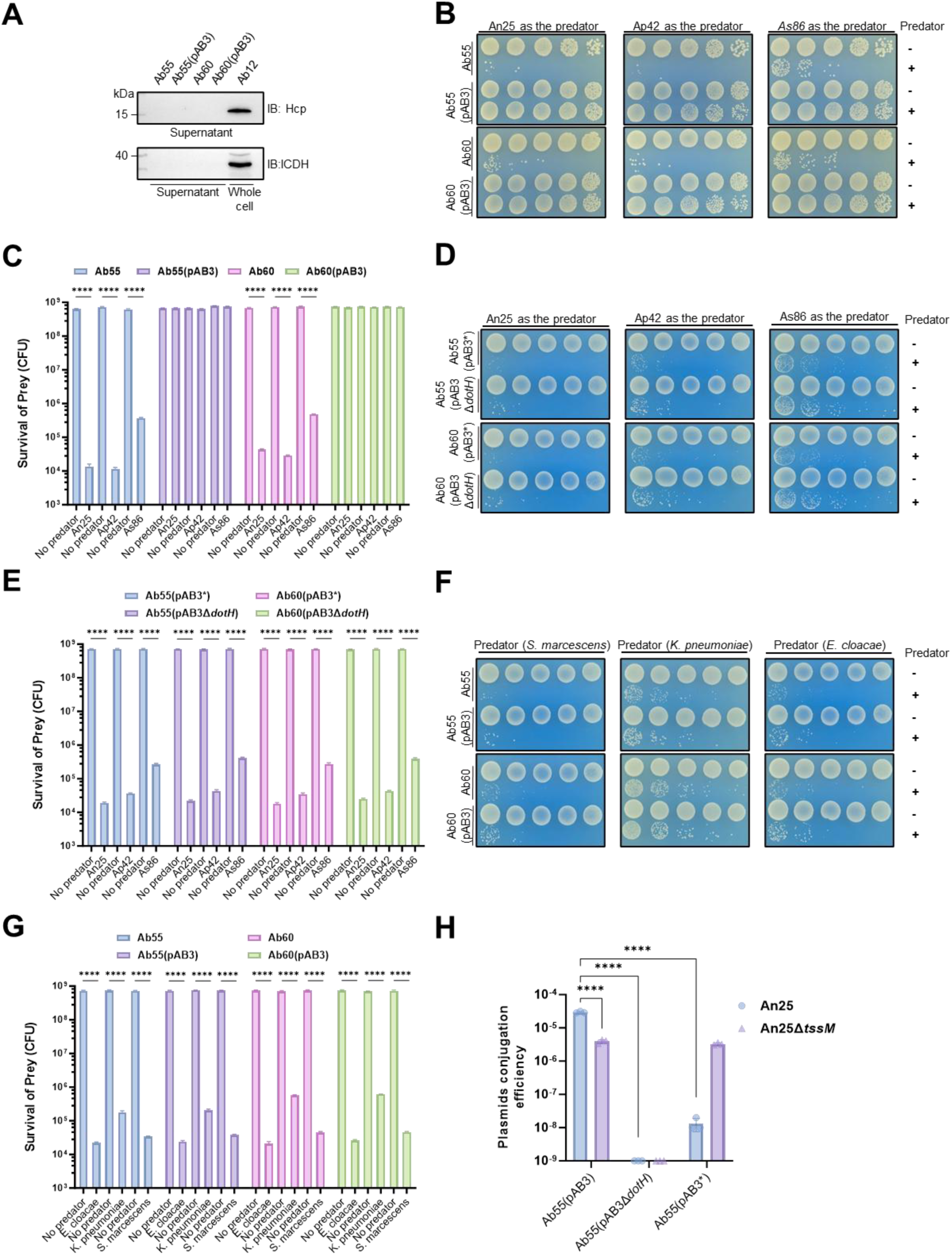
Conjugative dissemination of the pAB3 plasmid silences competitor T6SS to prevent fratricide *in vitro*. **(A)** Susceptibility of plasmidless *A. baumannii* to kin attacks. T6SS-active predators, including *A. nosocomialis* (An25), *A. pittii* (Ap42), and *A. seifertii* (As86), effectively eliminated the plasmidless, T6SS-defective *A. baumannii* prey strains Ab55 and Ab60. **(B-C)** Acquisition of wild-type pAB3 confers robust resistance against kin fratricide. Strains Ab55 and Ab60 harboring wild-type pAB3 were challenged by An25, Ap42, and As86. Representative images (B) and quantitative survival results (C) are shown. **(D-E)** Both T6SS repression and T4SS conjugation capabilities are essential for pAB3-mediated protection. Competition assays utilized Ab55 and Ab60 derivatives carrying either a T6SS-derepressed mutant (pAB3*) or a conjugation-defective mutant (pAB3Δ*dotH*). Neither mutant plasmid could protect the host from kin attacks. **(F-G)** pAB3 does not shield against distantly related pathogens. Strains Ab55(pAB3) and Ab60(pAB3) remained highly susceptible to *E. cloacae*, *K. pneumoniae*, and *S. marcescens*. **(H)** Interbacterial T6SS attacks actively stimulate pAB3 conjugation. *In vitro* conjugation assays were performed using Ab55 donors (harboring pAB3, pAB3*, or pAB3Δ*dotH*) and *A. nosocomialis* An25 or its Δ*tssM* mutant as recipients. Conjugation efficiency into the T6SS-competent An25 was significantly higher than into the T6SS-defective An25Δ*tssM*, indicating damage-responsive dissemination. For quantitative plots (C, E, G, H), data are presented as the mean ± SEM of three independent biological replicates. Statistical significance for CFU survival (C, E, G) and conjugation efficiencies (H) was determined using an unpaired, two-tailed Student’s *t*-test on log_10_-transformed values (****, *P* < 0.0001).

To dissect the mechanism of this pAB3-conferred resistance, we introduced two pAB3 derivatives into strains Ab55 and Ab60, respectively: a conjugation-defective mutant (pAB3Δ*dotH*) and a mutant that has lost the T6SS-repression activity (pAB3*). An earlier study found that two regions on pAB3 each is capable of inhibiting T6SS expression, one coding for two TetR-like transcriptional repressors and one harboring 8 predicted open reading frames mostly of unknown function ^21^. In interbacterial competition assays, the resulting strains carrying either pAB3* or pAB3Δ*dotH* remained completely sensitive to killing by strains An25, Ap42 or As86 (**Figure 5D-E**). As expected, because pAB3 cannot silence the T6SS of distantly related genera, its presence did not protect these strains from being killed by other Gram-negative pathogens such as *E. cloacae*, *K. pneumoniae*, and *S. marcescens* (**Figure 5F-G**). These results clearly reveal that a plasmid lacking the ability to suppress T6SS (pAB3*) or a plasmid incapable of conjugation (pAB3Δ*dotH*) has lost the ability to protect the host from T6SS-mediated killing by kin strains.

To evaluate the dynamics of this plasmid-mediated suppression during active interbacterial conflict, we performed *in vitro* conjugation assays using Ab55 harboring pAB3, pAB3ΔdotH or pAB3* as donors, and the *A. nosocomialis* strain An25 or its T6SS-defective mutant (An25Δ*tssM*) as recipients. Consistent with a previous report ^21^, the transfer efficiency of wild-type pAB3 into the T6SS-active An25 was significantly higher than that of the non-repressing pAB3* (**Figure 5H**). Such differences likely are caused by the killing of donor cells by the T6SS-potent recipient cells. As expected, pAB3Δ*dotH* failed to conjugate entirely due to the loss of the conjugation machinery. We also observed that when comparing different recipients, the conjugation efficiency of wild-type pAB3 into the T6SS-competent wild-type An25 was markedly higher than that of its transfer into the T6SS-defective An25Δ*tssM* (**Figure 5H**). This observation strongly implies that T6SS-mediated attack may actively stimulate or enhance the conjugative delivery of the silencing plasmid from the prey.

### Active plasmid transfer enables cooperative polymicrobial infection *in vivo*

We further investigated whether this plasmid-mediated T6SS suppression dictates infection outcomes in polymicrobial settings using the *Galleria mellonella* model ^22,23^. To track bacterial survival, we introduced the luminescence reporter pLuxCDABE into the Hcp-defective strain Ab55. While the T6SS-inactive Ab55 successfully established and maintained a systemic infection when co-inoculated with the T6SS-defective mutant An25Δ*tssM*, co-infection with the T6SS-active strain An25 led to rapid elimination of Ab55, evidenced by a drastic loss of bioluminescence signal (**Figure 6A**). In contrast, the introduction of wild-type pAB3 into Ab55(pLuxCDABE) completely rescued its survival, allowing it to successfully co-infect and persist together with An25 (**Figure 6A**). Measurement of the fluorescence intensity revealed that pAB3 maintained the *in vivo* bacterial load of the prey at levels comparable to the T6SS-defective co-infection control (**Figure 6B**). The introduction of pAB3* or pAB3Δ*dotH* failed to protect Ab55 from being eradicated by An25 (**Figure 6A-B**). These *in vivo* results corroborate well with our *in vitro* findings, demonstrating that an intact, conjugative T6SS-silencing plasmid is essential for the co-existence of different *Acinetobacter* strains in niches such as a host.

**Figure 6.**
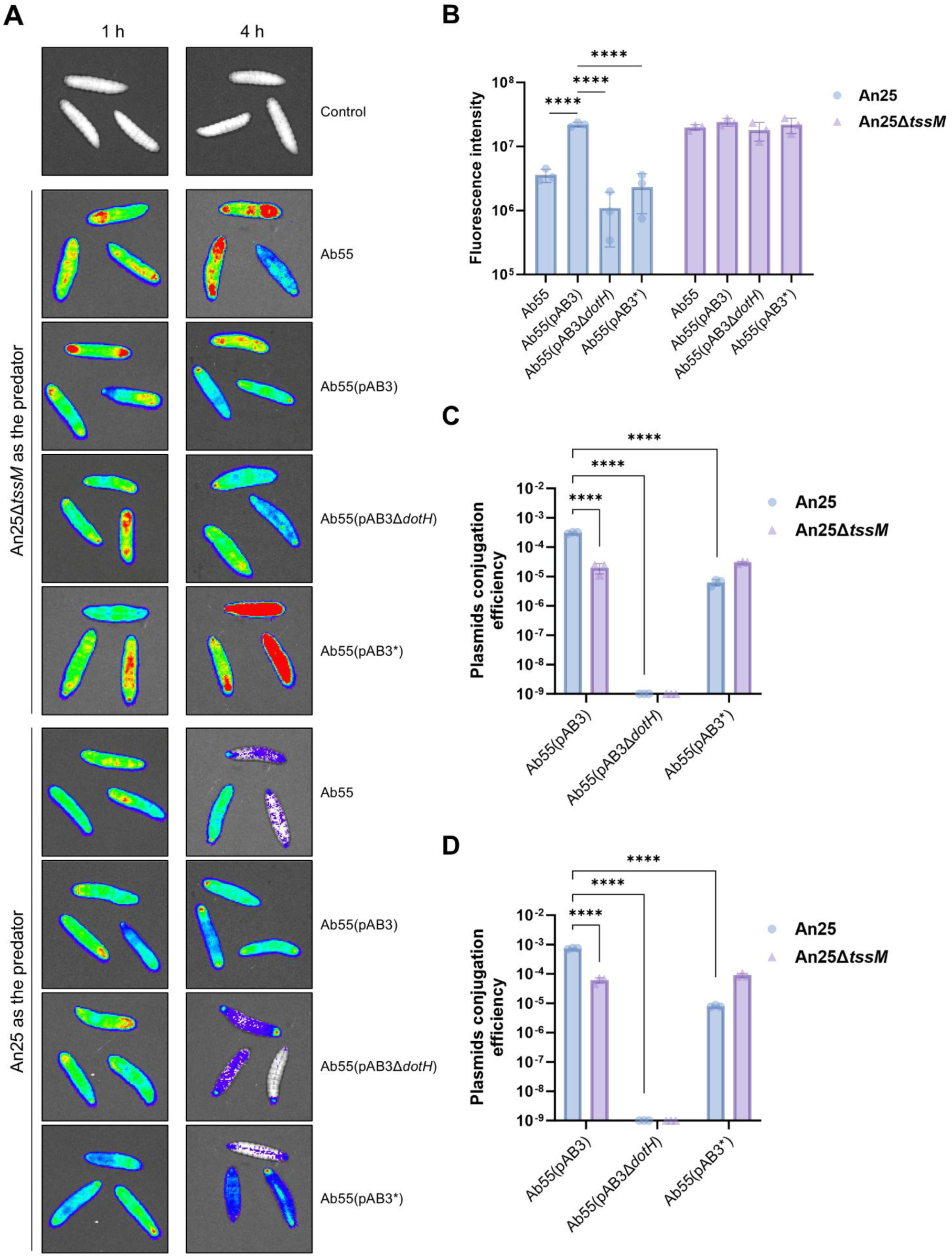
Active damage-responsive plasmid transfer enables cooperative polymicrobial infection *in vivo.* **(A-B)** pAB3-mediated suppression dictates *in vivo* infection outcomes. A luminescent derivative of strain Ab55 (pLuxCDABE) was co-inoculated into *G. mellonella* larvae with either the T6SS-active wild-type An25 or the defective An25Δ*tssM*. Ab55 prey strains harbored either no plasmid, wild-type pAB3, pAB3*, or pAB3Δ*dotH*. Representative bioluminescence images (A) and quantitative fluorescence intensities (B) demonstrate that acquisition of wild-type pAB3 uniquely rescued Ab55 from rapid *in vivo* elimination by An25. **(C)** Interbacterial antagonism enhances *in vivo* plasmid dissemination. Conjugation efficiencies from the Ab55 donor to An25 or An25Δ*tssM* recipients were assessed 6 hours post-co-inoculation in *G. mellonella*. Targeted transfer into the T6SS-active wild-type An25 was significantly higher than into the defective mutant. **(D)** Efficient plasmid transfer under sequential infection dynamics. *G. mellonella* larvae were first colonized by the recipient strains (An25 or An25Δ*tssM*), followed by inoculation of the Ab55 donor 2 hours later. Robust and targeted plasmid conjugation into the established wild-type An25 population was observed. Quantitative data in (B), (C), and (D) are presented as the mean ± SEM of three independent experiments. Statistical significance for fluorescence intensities (B) was determined using a one-way ANOVA followed by Tukey’s multiple comparisons test. Statistical significance for conjugation efficiencies (C, D) was determined using an unpaired, two-tailed Student’s *t*-test. All tests involving fluorescence or conjugation data were performed on log_10_-transformed values (****, *P* < 0.0001).

To directly confirm that plasmid sharing occurs dynamically during infection, we assessed the *in vivo* conjugation of pAB3 within the *G. mellonella* host. Co-inoculation of larvae with the Ab55 donor and the An25 recipient resulted in robust plasmid conjugation detectable 6 hours post-infection. Moreover, the *in vivo* conjugation efficiency was significantly higher than that observed when the same strains were mixed on the surface of LB agar (**Figure 6C**). This enhanced *in vivo* transfer aligns well with our earlier observation that interbacterial antagonism actively stimulates plasmid dissemination, as the confined and competitive niches within hosts likely forces extensive T6SS engagement.

To probe whether conjugation occurs between strains that cause sequential infections, we examined plasmid transfer under conditions where the donor strain was inoculated 2 hours after the recipient strain. Conjugation of pAB3 into the resident An25 population occurred efficiently (**Figure 6D**). Taken together, these findings demonstrate that plasmid-bearing strains secure their own survival by actively disseminating T6SS-silencing plasmid to aggressive peer bacteria, which silences the T6SS weapon and promotes polymicrobial infections.

## Discussion

In polymicrobial communities, such as those found in hospital-acquired pneumonias or wound infections, bacteria are engaged in a constant arms race for nutrient acquisition and niche dominance ^24,25^. Yet, strains lacking a functional T6SS also exhibit remarkable ecological resilience, frequently coexisting with T6SS-equipped competitors ^2-4,6^. Our findings reveal that the survival of *Acinetobacter* relies not only on a highly effective T6SS, but also on highly dynamic, multi-tiered defense systems, including physical shields and dissemination of T6SS-supression plasmids.

To gain an edge in competition against other T6SS-positive species, many *Acinetobacter* strains rely on a T6SS equipped with a highly effective effector arsenal, which subdues opponents prior to being overwhelmed (**Fig. 2**). This "offense as defense" strategy mirrors the ecological dynamic observed in the human gut, where commensal *E. cloacae* resist T6SS aggression from *Vibrio cholerae* by deploying a superior T6SS-mediated killing ability ^26^. For naturally T6SS-deficient strains, physical barriers appear to serve as the primary line of defense. Indeed, it has been shown that certain *K. pneumoniae* strains survive these identical T6SS attacks by creating a protective outer layer of type I CPS ^26^. Similarly, *V*. *cholerae* resist T6SS aggression by exopolysaccharide, preventing close contact with attacker cells ^27^. Recent studies indicate that in *A. baumannii* CPS acts as a double-edged sword, which can restrain its own T6SS deployment while simultaneously impeding T6SS assaults from competitors ^28^. Our analysis of the KL34 capsule reveals that the defensive efficacy of a capsule is not solely dictated by its thickness, but by its density. Despite being thinner than other capsule types, the high density of the KL34 capsule effectively prevents the penetration of T6SS from the competitor. This aligns with the observation that the high-density peptidoglycan cell wall of Gram-positive bacteria provides an effective physical barrier against most T6SS attacks ^29-31^.

Our transcriptomic data indicate that *Acinetobacter* does not passively endure T6SS strikes. Retaliatory offense has been documented in pathogens like *K. pneumoniae* ^32,33^; however, the response in *A. baumannii* appears to act by extensive gene expression reprogramming. Interbacterial assault induces not only the biosynthesis of the KL34 capsule and the PGA biofilm matrix, it also induces the expression of dormant T6SS structural genes (**Supplementary Tab. S1**). This transcriptomic shift aligns with the broader mechanism where bacteria activate stress response pathways (such as envelope stress responses) to repair damage caused by T6SS hits and fortify their cell envelope ^34^.

Our finding that the conjugative plasmid pAB3 functions as a vehicle to promote co-existence among *Acinetobacter* isolates unveils a new role for this enigmatic genetic element. The ability of pAB3-encoded TetR-like repressors to suppress T6SS expression has been well documented ^21,35^. It has been suggested that suppression of T6SS by this plasmid primarily serves to reduce metabolic burden, preventing the deleterious loss of resistance plasmids in non-antimicrobial environments ^17^. Our *in vivo* co-infection model offers a more ecologically sound explanation: active dissemination of this plasmid functions as a mechanism to prevent intra-species killing. Although the exact benefit awaits further investigation, co-existence of multiple *Acinetobacter* strains in a given niche may provide advantages such as cross-feeding and more effective in subduing host immunity.

Intriguingly, this plasmid-mediated attack-suppression is damage-responsive. We observed that the conjugation efficiency of pAB3 significantly escalates when the donor is subjected to an active T6SS attack (**Fig. 5H** and **Fig. 6C-D**), suggesting the existence of a damage-sensing regulatory mechanism. Recent findings show that exogenous peptidoglycan (PG) fragments, released during cell lysis or physical puncturing, act as universal danger signals that trigger defensive biofilm formation in various bacteria, including *A. baumannii* ^36^. Concurrently, the GacS/GacA two-component system in *A. baumannii* is known to coordinate central metabolism, virulence, and the Dot/Icm-like T4SS responsible for plasmid conjugation^20^. These observations together with our transcriptomic data suggest the following model: T6SS-induced membrane and PG damage serves as a danger signal that activates global stress networks (e.g., GacS/GacA), which also simultaneously fortifies the capsular shield, primes the retaliatory T6SS, and, crucially, hyper-activates the conjugative machinery to dispatch the silencing plasmid to the attacker.

This sophisticated defense network reflects an evolutionary trade-off. Maintaining a constitutively active T6SS is energetically costly and risks fratricide, which hinders population expansion. By utilizing a conjugative plasmid as a switch, *Acinetobacter* retains the potential to physically defend against attack from distantly related genera, while serving to promote a cooperative state with kins. The dynamic interplay of these defense mechanisms, including damage-responsive plasmid dissemination and capsular reprogramming, are often lost or attenuated in highly domesticated laboratory strains. By utilizing an array of recent clinical isolates, our study more accurately models the phenotypic diversity and interbacterial competitive dynamics associated with clinical infections. In clinical settings, where mixed-species or multi-strain infections are notoriously difficult to treat, the mutualism and antagonism mediated by machineries like the T6SS inevitably shape the pathogen composition. The plasmid’s role as an "ecological safeguard" may be a central driver behind the global success of multidrug-resistant *Acinetobacter* clones. Future strategies aiming at disrupting such cooperative compositions by targeting the T6SS or plasmid conjugation machineries could provide novel avenues for managing recalcitrant polymicrobial infections.

## MATERIALS AND METHODS

### Bacterial Strains and Growth Conditions

The bacterial strains used in this study are listed in Table S1. The majority of strains, including *A. baumannii*, *A. nosocomialis*, *A. pittii*, *A. seifertii*, *E. cloacae*, *S. marcescens*, and *K. pneumoniae*, were clinical isolates recovered from patients at the First Hospital of Jilin University, Changchun, China. All clinical isolates were verified by NGS. Unless otherwise specified, bacterial strains were routinely cultured at 37 °C in Luria–Bertani (LB) broth with shaking at 220 rpm, or on LB agar plates supplemented with 1.5% (wt/vol) agar. Antibiotics were added at the following concentrations when appropriate: ampicillin, 100 μg/mL; kanamycin, 30 μg/mL; gentamicin, 10 μg/mL; apramycin, 50 μg/mL; and streptomycin, 100 μg/mL.

### Clinical Cohort and Retrospective mNGS Data Collection

We retrospectively reviewed and analyzed clinical metagenomic next-generation sequencing (mNGS) diagnostic reports from 873 patients with suspected pulmonary infections at The First Hospital of Jilin University. This retrospective study was approved by the Institutional Review Board (IRB) and Ethics Committee of The First Hospital of Jilin University. Owing to the retrospective nature of the study using de-identified clinical data obtained from routine diagnostic reports, the requirement for written informed consent was formally waived by the Ethics Committee. All clinical investigations were conducted in strict accordance with the Declaration of Helsinki.

These mNGS assays were performed as routine clinical diagnostics by a College of American Pathologists (CAP)-accredited and ISO 15189-certified clinical laboratory. To strictly address the issue of potential background microbial DNA contamination (kit-ome) inherent to low-biomass samples (such as blood), the clinical testing pipeline implemented stringent quality control and diagnostic thresholding: (1) Negative controls (sterile water) were processed and sequenced in parallel with every batch of clinical samples to monitor and subtract environment and kit-derived contaminants; (2) A pathogen was classified as positive only if its sequence read count exceeded the established clinical diagnostic thresholds (stringent relative abundance or Reads Per Million [RPM] cut-offs calibrated for specific specimen types including blood, sputa, and BALF) and was confirmed by certified clinical microbiologists. Samples with suspected contamination or ambiguous clinical correlations were excluded from this study.

### DNA Manipulation and Plasmid Construction

Plasmids and primers used in this study are listed in Table S1. Bacterial genomic DNA was isolated using the TIANamp Bacteria DNA Kit (TIANGEN), and plasmid DNA was extracted using the TIANprep Mini Plasmid Kit (TIANGEN) according to the manufacturer’s instructions. PCR amplification was performed using TransStart FastPfu DNA Polymerase (TransGen Biotech). Unless otherwise noted, T4 DNA ligase and all restriction endonucleases were purchased from New England Biolabs (NEB).

Two expression plasmids were constructed and utilized in this study. pJL06, a derivative of the previously described pJL03 plasmid, drives constitutive gene expression under the control of an *Acinetobacter rpoD* promoter ^37^. This plasmid was used for gene complementation or ectopic expression in *Acinetobacter* species where indicated. Additionally, pZY01, derived from the commercial plasmid pUC59-kana, carries a tac promoter to enable efficient gene expression in *Enterobacteriaceae* species, including *E. cloacae*, *S. marcescens*, and *K. pneumoniae*. Genes of interest (e.g., the tetR-family repressors and the KL34 capsule locus genes) were amplified and cloned into the corresponding vectors using BamHI/SalI or XbaI/NotI restriction sites. The resulting constructs were introduced into their respective host strains for subsequent protein production and phenotypic analyses.

### Bacterial Transformation

Electroporation was used to introduce plasmids into *Acinetobacter* and *Enterobacteriaceae* species. Briefly, overnight cultures originating from a single colony were diluted 1:100 into fresh LB broth and grown to the mid-logarithmic phase. Cells were harvested at 4 °C, washed three times with ice-cold 10% (vol/vol) glycerol, and resuspended in the same solution. Aliquots were prepared and stored at −80 °C until use. For electroporation, 1 μg of plasmid DNA was mixed with 100 μL of electrocompetent cells and subjected to a single pulse (25 μF, 2.5 kV/cm, 200 Ω) using a Gene Pulser Xcell system (Bio-Rad). Immediately after pulsing, cells were recovered in pre-warmed LB broth at 37 °C for 1 h and subsequently plated on LB agar supplemented with the appropriate selective antibiotic. Positive transformants were verified by colony PCR.

Triparental mating was employed for strains refractory to electroporation. Donor, recipient, and helper strains (*E. coli* MT607 carrying pRK6001 ^38^) were mixed at equal ratios, pelleted by centrifugation, and washed three times with fresh LB broth. The cell mixture was spotted onto LB agar and incubated at 37 °C for 4–6 h. Following incubation, cells were resuspended and plated onto selective media to isolate transconjugants.

### Construction of Bacterial Mutants

Markerless gene deletions were generated via homologous recombination using the R6Kγ-based suicide allelic exchange vector pSR47s. Approximately 1-kb DNA fragments flanking the upstream and downstream regions of each target gene were amplified from genomic DNA, fused by overlap extension PCR, and cloned into pSR47s digested with SacI and SalI. This allelic replacement strategy substitutes the target gene with an open reading frame encoding a 30-amino-acid scar peptide composed of the first 15 and the last 15 residues of the native protein.

The constructed knockout plasmids were introduced into recipient strains via triparental mating using *E. coli* MT607 harboring pRK6001 as the helper strain. Following conjugation, transconjugants were selected and subsequently cultured on LB agar plates supplemented with 5% (wt/vol) sucrose. This step counter-selects against the sacB-containing suicide vector, thereby enriching for cells that had undergone a second homologous recombination event. Deletion mutants were identified by colony PCR using primer pairs annealing outside the homologous arms and were further verified by Sanger sequencing to confirm correct allelic exchange.

### Antibodies and Immunoblotting

Standard immunoblotting procedures were performed using the following primary antibodies: rabbit anti-Hcp (1:2,000), rabbit anti-isocitrate dehydrogenase (ICDH) (1:2,000), mouse anti-Flag (1:3,000; Sigma, F1804), and mouse anti-HA (1:3,000; Sigma, H3663). Crucially, to accurately detect Hcp across phylogenetically distinct genera and prevent cross-reactivity artifacts, we utilized two distinct primary antibodies. For the *Enterobacteriaceae* species (*K. pneumoniae*, *E. cloacae*, and *S. marcescens*), we employed a broadly cross-reactive rabbit polyclonal anti-Hcp antibody originally raised against *Klebsiella* Hcp, which leverages the high structural homology among these related enteric species. Conversely, for all *Acinetobacter* isolates, a custom rabbit polyclonal antibody specifically raised against the *A. baumannii* Hcp peptide sequence was generated and utilized to ensure high specificity. To analyze secreted proteins, bacterial cultures were first centrifuged at 4,000 × *g* for 5 min to remove cells. The resulting supernatants were collected and further clarified by centrifugation at 12,000 × *g* for 1 min. A 40-μL aliquot of each clarified supernatant was mixed with 10 μL of 5× SDS sample buffer, boiled for 10 min, and centrifuged at 12,000 × *g* for 1 min. The resulting supernatants were used for SDS-PAGE analysis. For the analysis of whole-cell samples, bacterial pellets corresponding to an optical density at 600 nm (OD600) of 1.0 (approximately 1 × 10^9^ cells) were directly resuspended in 50 μL of 1× SDS sample buffer, boiled for 10 min, and centrifuged at 12,000 × *g* for 3 min.

Protein samples were resolved by SDS-PAGE, transferred onto nitrocellulose membranes, and blocked with 5% (wt/vol) nonfat milk (Bio-Rad) for 1 h at room temperature. Membranes were incubated with primary antibodies overnight at 4 °C. After three washes with PBST, the membranes were incubated with appropriate IRDye-labeled or horseradish peroxidase (HRP)-conjugated secondary antibodies for 1 h at room temperature. Signals were visualized using an Odyssey CLx imaging system (LI-COR) or a ChemiDoc MP system (Bio-Rad).

### Interspecies Bacterial Competition Assay

Intra- and interspecies T6SS-mediated killing assays were performed as previously described ^9^. Briefly, predator and prey strains from overnight cultures were washed with sterile phosphate-buffered saline (PBS), resuspended in fresh LB broth, and normalized to an identical OD600. Predator strains were mixed with prey strains carrying appropriate natural or plasmid-encoded antibiotic resistance markers at a 1:1 ratio. The mixtures were centrifuged briefly, and the pellets were resuspended in 20 μL of fresh LB broth. The cell mixtures were spotted onto LB agar plates and incubated at 37 °C for 4–6 h. Bacterial spots were then harvested from the agar surface, resuspended in sterile PBS, serially diluted, and plated onto selective LB agar to enumerate surviving prey cells. Prey strains processed in parallel without predator exposure served as initial viable cell count controls.

### RNA Extraction and qPCR Analysis

Total bacterial RNA was extracted using the MolPure Bacterial RNA Kit (Yeasen) according to the manufacturer’s instructions. Residual genomic DNA was eliminated, and complementary DNA (cDNA) was synthesized using the PrimeScript RT Reagent Kit (Takara). Quantitative real-time PCR (qPCR) was performed on a CFX Connect Real-Time PCR Detection System (Bio-Rad) using BlasTaq 2× qPCR MasterMix (ABM). Each reaction contained 100 ng of cDNA and 0.4 μM of gene-specific primers (listed in Table S1). Expression levels were normalized to the internal reference gene rpoC. Relative gene expression was calculated using the standard 2^−ΔΔCT^ method.

### TEM Sample Preparation and Observation

*A. baumannii* cells from overnight cultures were subcultured at a 1:100 dilution and grown to the mid-logarithmic phase. Cells were harvested by centrifugation (4,000 × *g*, 20 min, 4 °C) and immediately fixed in 4% (wt/vol) glutaraldehyde prepared in 0.1 M phosphate buffer (Macklin) for 24 h at 4 °C in the dark. Samples were washed thoroughly and post-fixed with 1% (wt/vol) osmium tetroxide (OsO_4_) for 2 h at room temperature in the dark. Samples were then washed twice with double-distilled water (ddH_2_O) for 10 min each and dehydrated through a graded ethanol and acetone series. Dehydrated cells were infiltrated with a 1:1 mixture of propylene oxide and epoxy resin for 1–2 h, followed by pure resin for an additional 1–2 h. Samples were embedded in molds and polymerized overnight at 35 °C. Ultrathin sections (50–70 nm) were prepared using an EM UC7 ultramicrotome (Leica), collected on copper grids, and sequentially stained with uranyl acetate and lead citrate. Samples were visualized using an HT7800 transmission electron microscope (Hitachi).

For the quantitative analysis of capsular thickness and electron density, TEM micrographs were analyzed using ImageJ software (NIH). To determine capsular thickness, the spatial scale was first globally calibrated against the scale bar provided in each original micrograph. The Straight-Line tool was then utilized to measure the distance from the outer membrane to the outer capsular boundary at multiple random locations per bacterial cell.

### Plasmid Conjugation Assay

The transfer efficiency of plasmid pAB3 and its derivatives among *Acinetobacter* species was evaluated as previously described ^39^. An apramycin resistance cassette was integrated into pAB3 via homologous recombination. *A. baumannii* strain Ab55 derivatives (kanamycin-sensitive) harboring wild-type pAB3, a *dotH*-deficient mutant (pAB3Δ*dotH*), or a T6SS-derepressed mutant (pAB3*) were used as donors. The kanamycin-resistant, apramycin-sensitive *A. nosocomialis* strain An25 or its T6SS-defective mutant (An25Δ*tssM*) served as the recipient.

For *in vitro* assays, washed donor and recipient cells were mixed at a 1:5 ratio in LB broth, spotted onto LB agar, and incubated at 37 °C for 6 h. Transconjugants were quantified by plating on dual-selective media. For *in vivo* conjugation assays, *G. mellonella* larvae were infected either simultaneously (donor and recipient injected together at a 1:5 ratio) or sequentially (the recipient strain was inoculated first to establish colonization, followed by the donor strain 2 h later). Following a 6-h incubation at 37 °C, the bacterial populations were recovered from the larval hemocoel and plated on selective media to enumerate transconjugants. As a negative control, homogenates from uninfected larvae were plated on the same selective media, which yielded no bacterial growth, thereby confirming the absence of endogenous flora capable of growing under these selective conditions.

### *G. mellonella* Infection Model and *In Vivo* Imaging

To track donor bacteria in real time, the bioluminescent reporter plasmid pJL06::luxCDABE was constructed. This self-sufficient operon encodes both luciferase and its endogenous substrate biosynthesis machinery, allowing continuous bioluminescence without exogenous substrates. The plasmid was introduced into *A. baumannii* donor strains. *G. mellonella* larvae were inoculated with mixed bacterial suspensions (10 μL per larva; ∼1×10^6^ CFU total). Bacterial survival, colonization, and infection dynamics were continuously monitored *in vivo* using an IVIS Lumina III imaging system (Revvity).

### Genome Assembly, Quality Control, and Genomic Annotation

*De novo* assembly of clinical bacterial genomes was performed using SPAdes (v3.12.0) ^40^ with default parameters. Species identification was conducted using pubMLST ^41^ and phyloFlash (v3.4.2) ^42^. Genome quality was assessed via CheckM (v1.2.2) ^43^, retaining only assemblies with completeness > 98% and contamination < 5%. Contaminated or mixed genomes were binned using MetaBAT2 (v2.15) ^44^, and valid bins were retained. Ultimately, 190 high-quality clinical *A. baumannii* isolates were selected.

Publicly available whole-genome data were retrieved from the NCBI database. Stringent quality filtering (CheckM: completeness > 99%, contamination < 1%, contigs < 200, N50 > 20,000) yielded 6,710 *Acinetobacter* spp., 4,805 *Enterobacter* spp. (528 *E. cloacae*), 4,387 *Neisseria* spp. (37 *N. subflava*), and 2,171 *Serratia* spp. (1,612 S. marcescens) genomes. Because *Klebsiella* spp. and *Pseudomonas* spp. were markedly overrepresented in terms of genome availability compared with other taxa, a stratified random sampling approach based on MLST (v2.23.0) ^45^ sequence types (STs) was employed, randomly selecting a 10% representative subset. This yielded 3,982 *Klebsiella* spp. (3,605 *K. pneumoniae*) and 8,141 *Pseudomonas* spp. (3,287 *P. aeruginosa*) genomes after quality control. All retained genomes were uniformly annotated using Prokka (v1.11) ^46^. A schematic overview of the complete bioinformatic workflow and data processing pipeline is illustrated in **Figure S11**.

### Identification of T6SS Loci and VgrG Effectors

Experimentally validated T6SS core genes from the SecReT6 database were used for homology-based clustering via Proteinortho (v6.0.29) ^47^. Contiguous genomic regions containing 11 T6SS core genes in *Acinetobacter* (lacking *tssJ*) and *Klebsiella* were classified as complete loci (T6SS+). Genus−specific completeness thresholds were applied (≥12 core genes for *Pseudomonas*, *Serratia*, *Enterobacter*, and *Neisseria*). The *tssI* (*vgrG*) gene was excluded from locus typing due to its highly variable copy number and genomic position.

A Hidden Markov Model (HMM) profile for VgrG was constructed using HMMER (v3.2) ^48^ (E-value < 1×10^−6^). The relative abundance of *vgrG* per genome was normalized against the number of core genes (using *tssB* as reference). Rank-sum tests were performed to compare the distribution of average *vgrG* copy numbers across genera.

### Typing of Offensive T6SS Loci

Based on established literature defining T6SS loci associated with interbacterial antagonism, we mapped the proportion of "offensive" T6SS loci across species. In *A. baumannii*, this locus corresponds to the conserved gene arrangement: *tssB*, *tssC*, *tssD, tssE, tssF, tssG, tssM, tssH, tssA, tssK, tssL*. Similarly specific core gene arrangements were used to define the offensive H1-T6SS in *P. aeruginosa* ^49^, as well as corresponding offensive loci in *E. cloacae* ^50^, *K. pneumoniae* ^33^, and *S. marcescens* ^51^. For *N. subflava*, where functional differentiation is uncharacterized, all complete T6SS loci were included in the analysis.

### Comparative Genomics, KL Typing, and Phylogenetic Analysis

Comparative genomics was performed on 44 clinical *A. baumannii* isolates, comprising 6 T6SS-resistant and 38 T6SS-sensitive isolates. Genes shared exclusively among insensitive isolates were identified using CD-HIT (v4.8.1) ^52^ (60% sequence identity and 60% length coverage thresholds). The KL types of 190 clinical *A. baumannii* strains were typed using KAPTIVE (v3.0.0b5) ^53^, and specific KL components were mapped via BLASTp (v2.12.0+)^54^. Genomic loci were visualized using ChiPlot. A maximum-likelihood phylogenetic tree of 62 representative genomes was inferred from core marker genes (PhyloPhlAn v3.0.67) ^55^ using IQ-TREE (v2.1.4) ^56^ with 10,000 ultrafast bootstraps, and visualized using iTOL (v7.1) ^57^.

### Transcriptomic Analysis

Total RNA from *A. baumannii* Ab27 co-cultured with *E. cloacae* or its Δ*tssM* mutant was sequenced on an Illumina platform. Initial quality assessment of the raw sequencing data confirmed excellent metrics, with over 93% of the bases achieving a Phred quality score of ≥ 30 (Q30 > 93%). Raw reads were subsequently subjected to rigorous quality control using fastp ^58^. Specifically, reads were completely discarded if they contained adapter sequences, ambiguous bases (N), or if more than 5% of the bases within a given read possessed a Phred quality score of ≤ 5. The resulting high-quality clean reads were mapped to the *A. baumannii* reference genome using Bowtie2 (v2.2.3) ^59^. To ensure absolute specificity and eliminate false positives during downstream quantification, the alignments were subjected to strict post-mapping filtration: reads with a mapping quality (MAPQ) score of < 10, non-concordantly aligned read pairs, and multi-mapped reads were entirely excluded. Transcriptome assembly and operon identification were performed with Rockhopper ^60^, and highly confident gene expression quantification was executed using HTSeq (v0.6.1) ^61^. Differential expression analysis was conducted using the DESeq2 R package (v1.20.0) ^62^. Statistical significance was rigorously defined by an adjusted P-value < 0.05. Given the extreme purity of our stringently filtered dataset, a fold-change cutoff of log2(fold change) ≥ 0.3 was strategically applied. This highly sensitive threshold was optimized to reliably capture the coordinated, operon-wide expression shifts within specific defensive gene clusters (e.g., the massive KL34 capsule biosynthesis locus) that were a priori identified via our comparative genomics analysis between T6SS-resistant and T6SS-sensitive strains.

### Prediction of TetR-Family Transcription Factor Binding Sites

The operon architectures of the extracted T6SS loci were predicted using Operon-mapper ^63^. Bacterial promoter elements (−10 and −35 boxes) within the 150-bp upstream regulatory regions were identified using BPROM. Palindromic binding motifs were predicted using MEME (v5.5.7) ^64^ with motif widths constrained between 12 and 20 bp. Multiple sequence alignments of these promoter regions were generated using MAFFT and visualized in Jalview (v2.11.4.1) ^65^.

### Statistical Analysis

All statistical analyses and data visualization were performed using GraphPad Prism software (version 10.0). Quantitative data derived from at least three independent biological replicates (n≥3) are presented as the mean ± standard error of the mean (SEM). To satisfy the assumptions of parametric tests (normal distribution and homogeneity of variance), bacterial survival counts (CFU), plasmid conjugation frequencies, and fluorescence intensities spanning multiple orders of magnitude were log10-transformed prior to statistical evaluation. Differences between two independent experimental groups were analyzed using an unpaired, two-tailed Student’s t-test. Comparisons among three or more groups were evaluated using a one-way analysis of variance (ANOVA) followed by Dunnett’s multiple comparisons test (when comparing various experimental groups to a single control) or Tukey’s multiple comparisons test (for all-pairwise comparisons). For relative gene expression (RT-qPCR), unpaired Student’s t-tests were utilized. For capsular thickness and electron density measurements, data from multiple individual bacterial cells (n≥15) were analyzed using a one-way ANOVA followed by Tukey’s multiple comparisons test. A P-value of < 0.05 was considered statistically significant (* P < 0.05; ** P < 0.01; *** P < 0.001; **** P < 0.0001; ns, not significant).

## Acknowledgements

This study was funded in part by Jilin Science and Technology Agency grant 20260205009GH (LS).

## Author contributions

L.S. directed and conceptualized the study. Experimental procedures were executed by J.J., X.D., S.G., and X.Z., while bioinformatic analyses were performed by M.Z. and Q.G. Data interpretation and analysis were conducted by L.S., D.L., and Z.Q.L. L.S. drafted the initial manuscript. L.S. and Z.Q.L. revised the manuscript, with critical input and final approval from all authors.

## Figures and Legends

**Figure S1.**
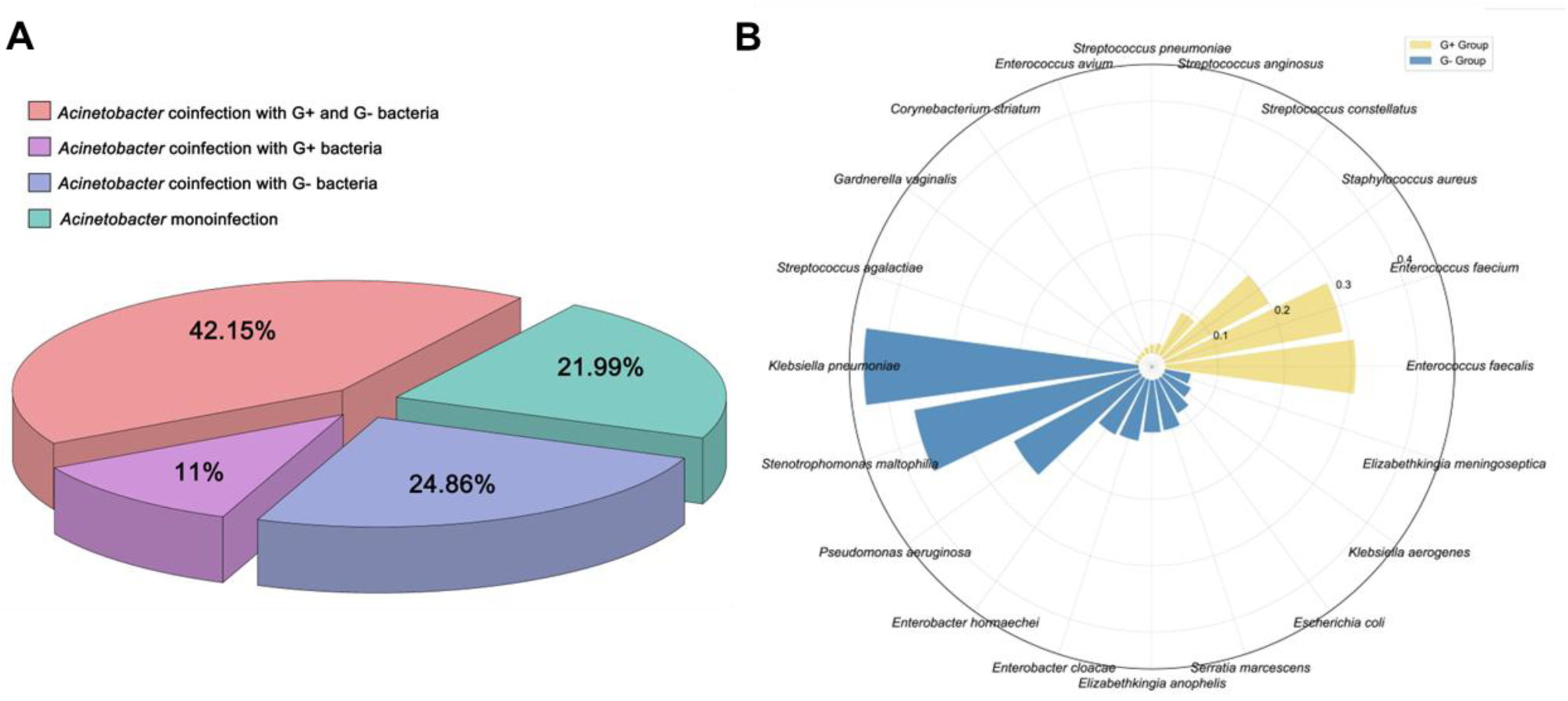
Pathogenic *Acinetobacter* spp. are highly prevalent in polymicrobial clinical infections. The microbial composition of 873 clinical specimens (blood, sputa, and BALF) collected from patients with pulmonary infections was determined by NGS. **(A)** The pie chart illustrates the proportion of samples representing mono-infections caused solely by *Acinetobacter* spp. (21.99%), as opposed to polymicrobial infections (78.01%). The polymicrobial cases are further subdivided into coinfections involving Gram-positive bacteria (11.00%), Gram-negative bacteria (24.86%), or a complex consortium of both (42.15%). **(B)** Among these polymicrobial cases, the most frequently co-occurring bacterial pathogens alongside Acinetobacter spp. are *K. pneumoniae* (41.43%), *S. maltophilia* (34.41%), and *E. faecalis* (28.70%).

**Figure S2.**
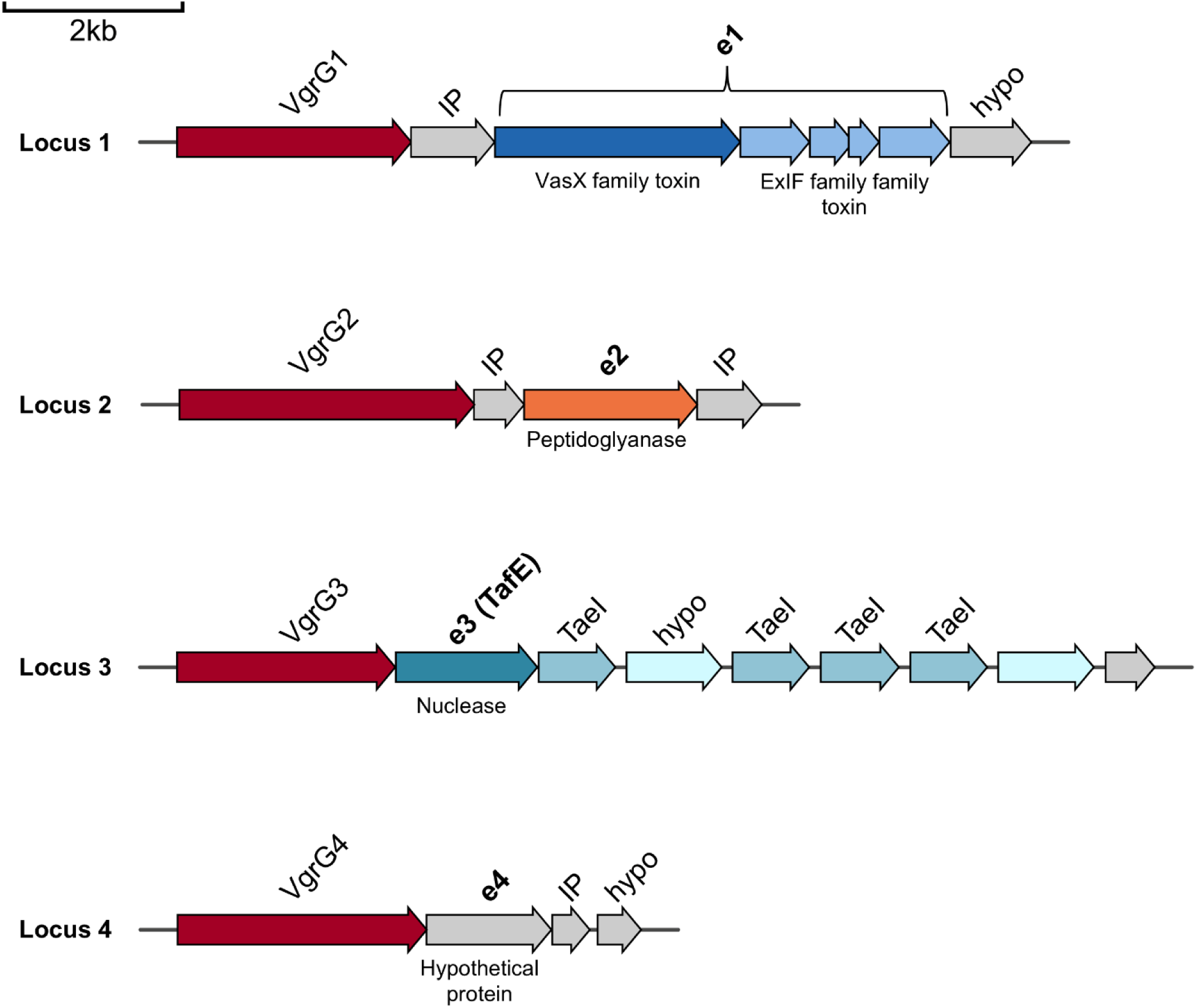
Organization of VgrG-associated effector loci in *A. baumannii* strain Ab12. Schematic representation of the four *vgrG* gene clusters identified in the genome of the clinical isolate Ab12. The predicted effector genes downstream of each *vgrG* are designated as *e1* to *e4*. Locus 1 encodes a cluster of five potential effectors (collectively referred to as *e1*), comprising one protein with a VasX_N domain and four proteins containing Burk_ExIF domains. Locus 2 encodes a putative peptidoglycanase effector carrying a LysM domain (*e2*). Locus 3 encodes the DNase effector TafE (*e3*). Locus 4 harbors a downstream gene coding for a protein of unknown function (*e4*). Gray arrows indicate putative immunity proteins or hypothetical proteins. Scale bar: 2 kb.

**Figure S3.**
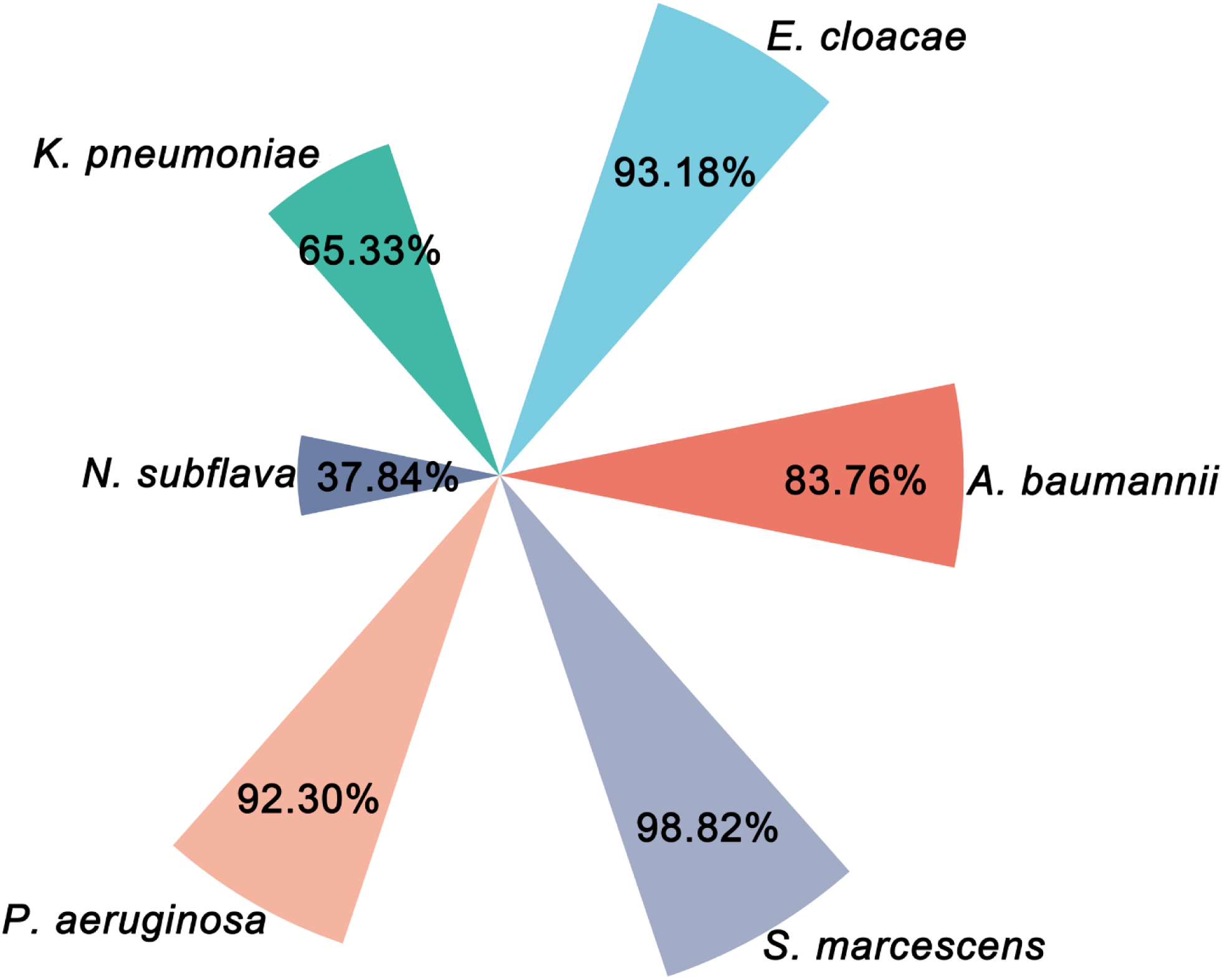
Distribution of intact T6SS loci across genomes of *A. baumannii* and other Gram-negative bacteria. The frequency of genomes encoding a complete T6SS locus was analyzed using public genomic databases. The chart illustrates the percentage of T6SS-positive isolates for *S. marcescens* (98.82%), *E. cloacae* (93.18%), *P. aeruginosa* (92.30%), *A. baumannii* (83.76%), *K. pneumoniae* (65.33%), and *N. subflava* (37.84%). Notably, approximately 16.2% of *A. baumannii* genomes lack a complete T6SS cluster.

**Figure S4.**
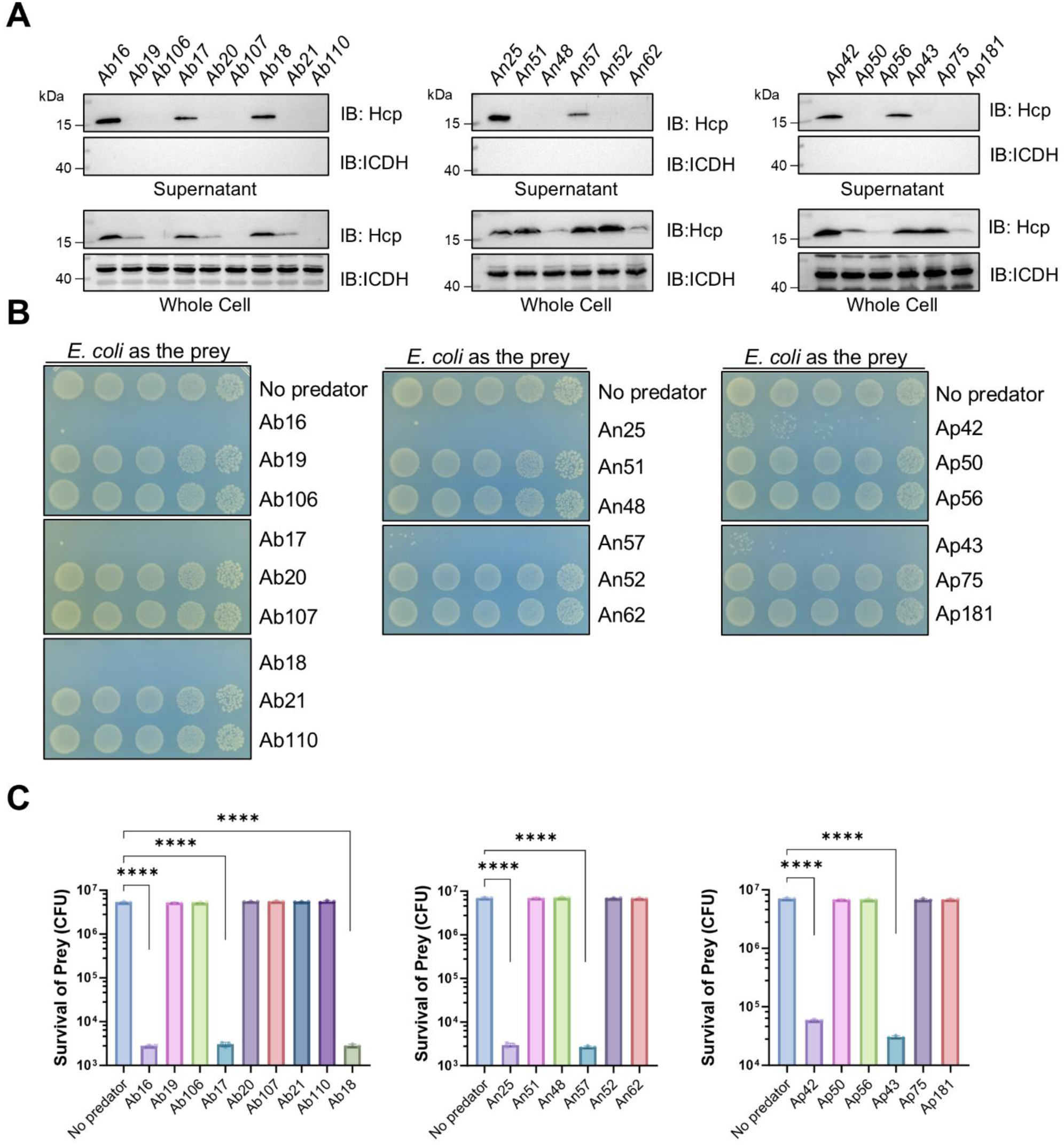
Survey of T6SS activity among clinical *Acinetobacter* isolates. **(A)** Hcp secretion in representative clinical isolates of *A. baumannii* (Ab), *A. nosocomialis* (An), and *A. pittii* (Ap). Among the 186 strains tested, only those actively secreting Hcp into the supernatant were considered T6SS-positive. ICDH was probed as a loading control for whole-cell lysates. **(B-C)** Functional assay of T6SS activity against *E. coli*. Representative images (B) and quantitative survival results (C) demonstrate that isolates exhibiting active Hcp secretion consistently executed T6SS-dependent killing of *E. coli*. For (C), quantitative data are presented as the mean ± SEM of three independent biological replicates performed in triplicate. Statistical significance was determined using an unpaired, two-tailed Student’s *t*-test on log_10_-transformed CFU values (****, *P* < 0.0001).

**Figure S5.**
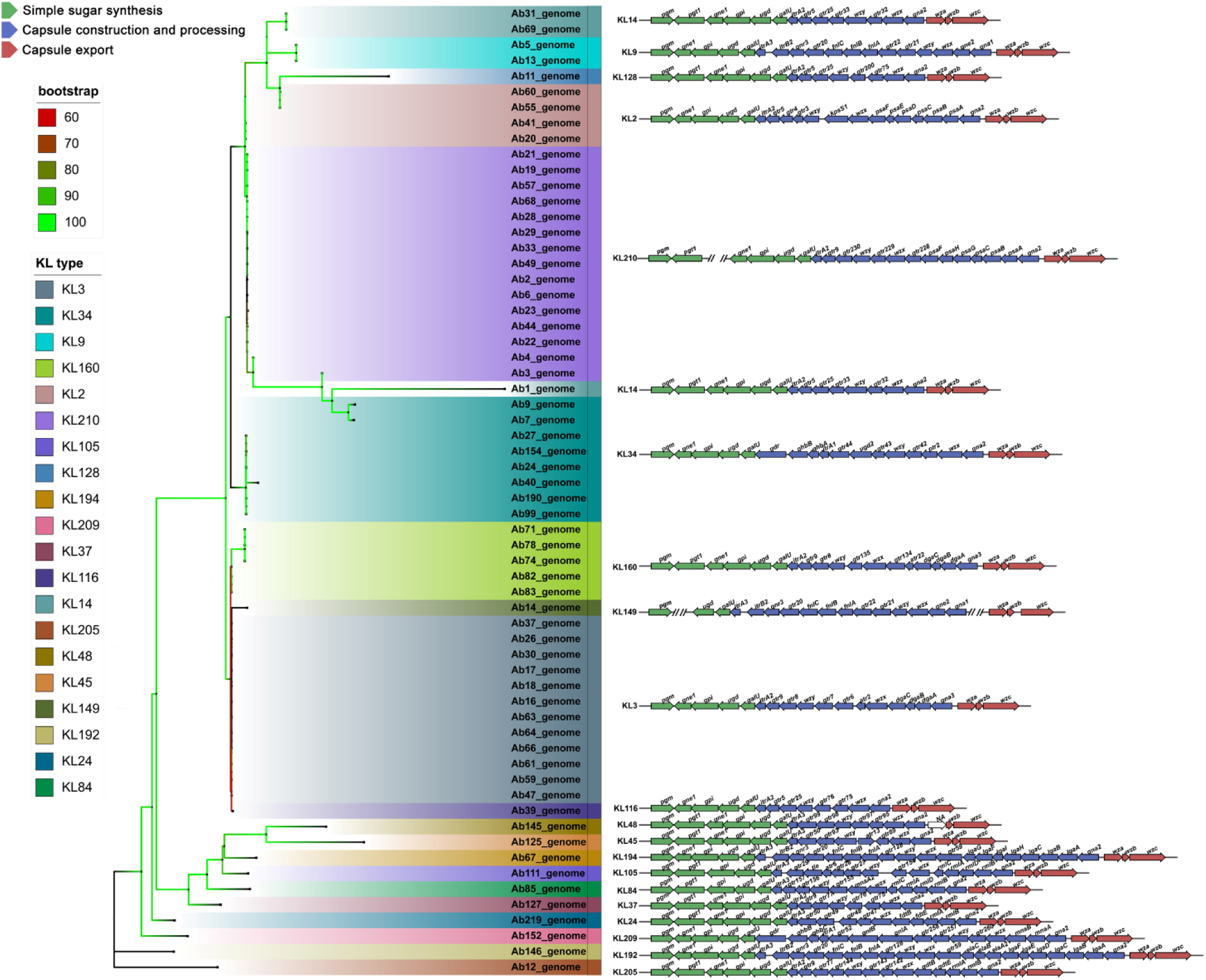
Phylogenetic distribution and genetic organization of capsule biosynthesis loci in clinical *Acinetobacter* isolates. A maximum-likelihood phylogenetic tree illustrates the evolutionary relationships among the *Acinetobacter* clinical isolates. Colored backgrounds group strains by their specific capsular polysaccharide (KL) locus type. Notably, all T6SS-resistant strains (e.g., Ab27, Ab154, Ab24, Ab40) cluster exclusively within the KL34 lineage. The right panel displays the linear genetic organization of representative capsule loci, with genes color-coded by predicted function: simple sugar synthesis (green), capsule construction and processing (blue), and capsule export (red).

**Figure S6.**
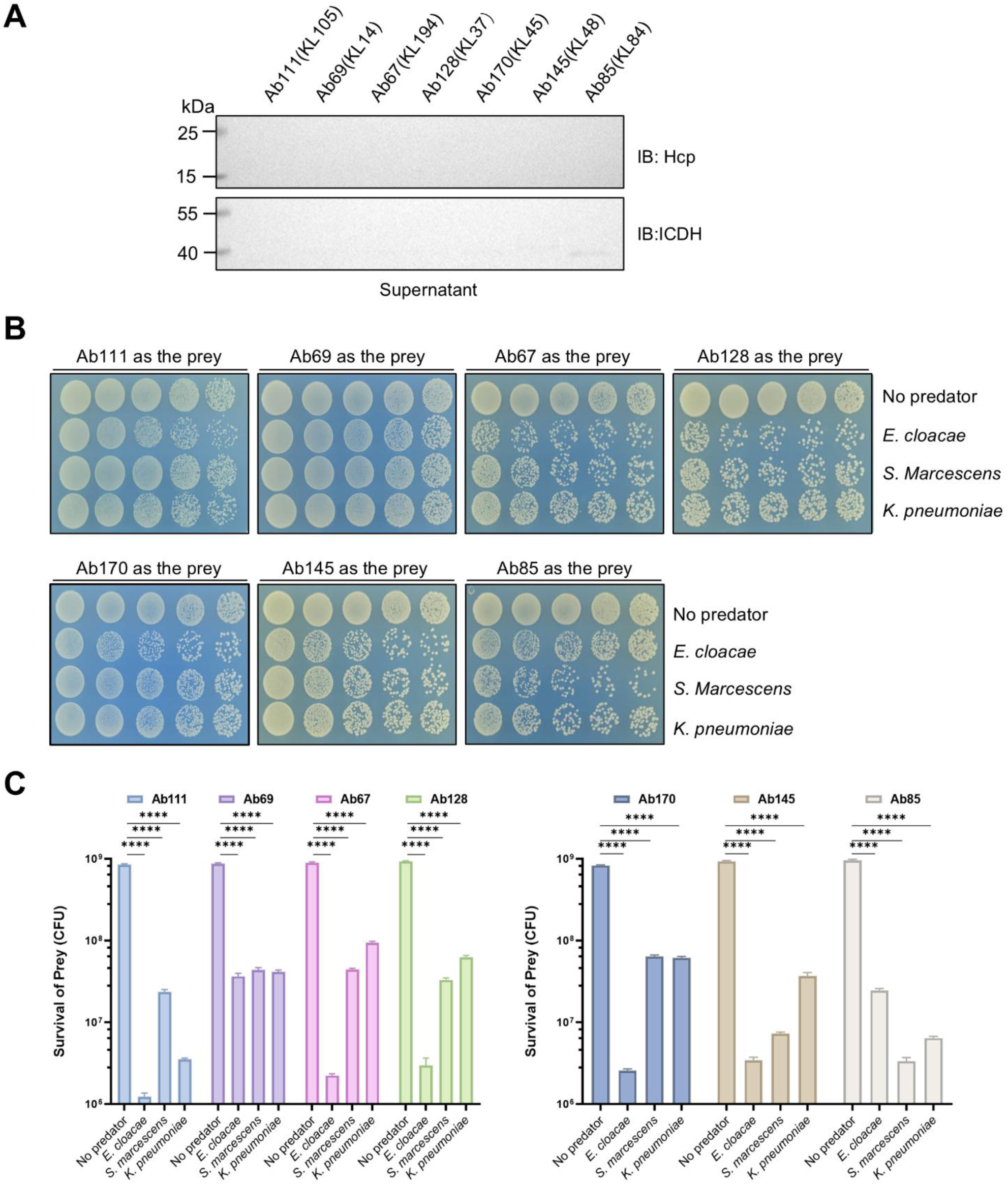
T6SS-deficient *A. baumannii* strains harboring non-KL34 capsule types are highly susceptible to interbacterial killing. **(A)** Hcp secretion profiles of clinical *A. baumannii* isolates representing various non-KL34 capsule types (KL105, KL14, KL194, KL37, KL45, KL48, and KL84). **(B-C)** Killing of T6SS-defective *A. baumannii* isolates by Gram-negative pathogens. Representative images (B) and quantitative survival results (C) are shown. Unlike the highly resistant KL34-type strains, all non-KL34 isolates were susceptible to predator attacks. For (C), data are presented as the mean ± SEM of three independent biological replicates performed in triplicate. Statistical significance was determined using an unpaired, two-tailed Student’s *t*-test on log_10_-transformed CFU values (****, *P* < 0.0001).

**Figure S7.**
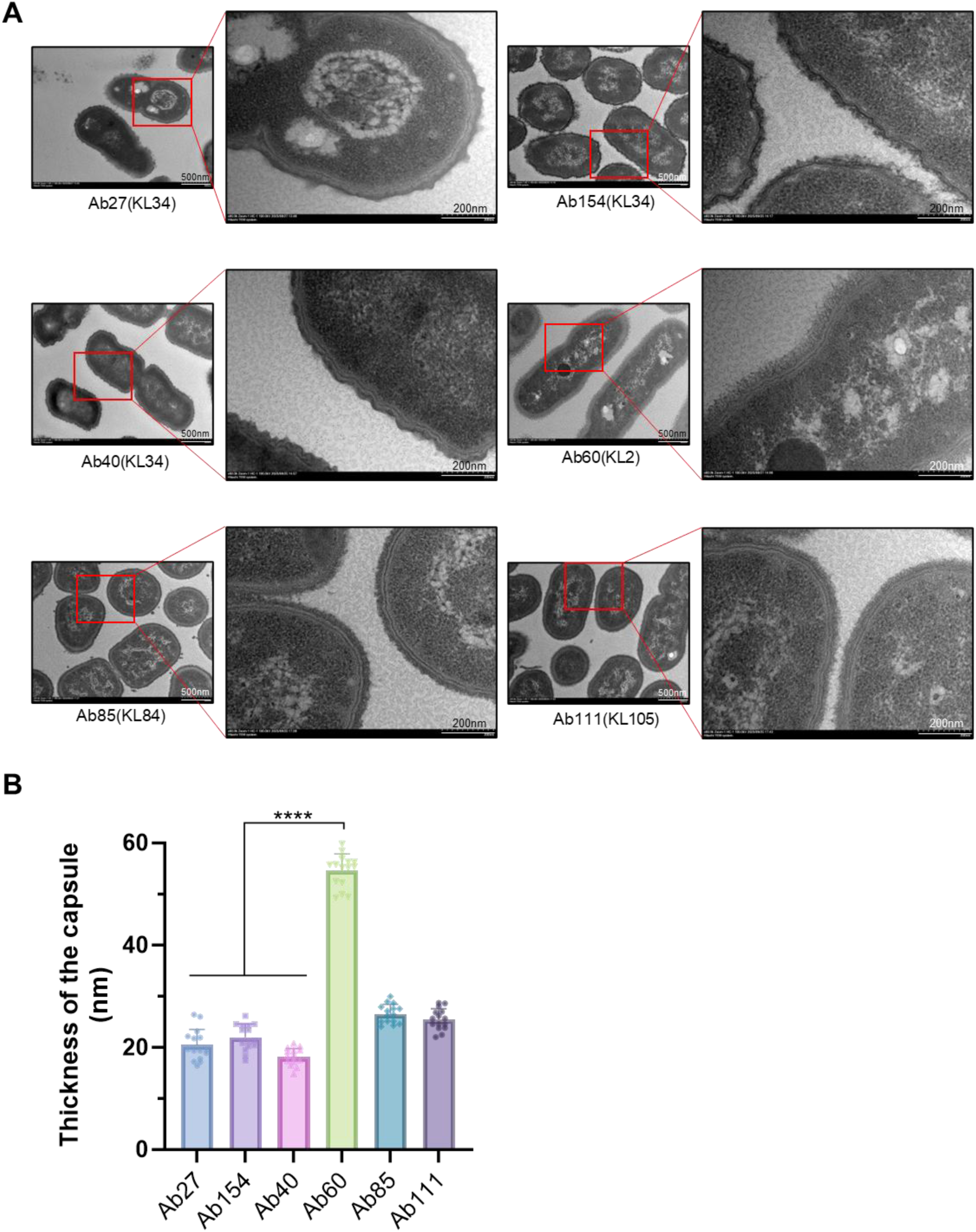
The KL34-type capsule exhibits exceptionally high electron density. **(A)** TEM visualization of the capsular layers in T6SS-resistant (KL34-type: Ab27, Ab154, Ab40) and T6SS-sensitive (KL2-type: Ab60; KL84-type: Ab85; KL105-type: Ab111) *A. baumannii* strains. Red boxes indicate the magnified regions (right/inset panels). Scale bars: 500 nm (low magnification) or 200 nm (high magnification). **(B)** Quantification of capsule thickness measured by ImageJ. Notably, the KL34 capsule is quantitatively thinner than the KL2-type capsule of Ab60, and shows no statistically significant difference compared to Ab85 (KL84) or Ab111 (KL105). Data are presented as the mean ± SEM of multiple individual bacterial cells (n ≥ 15 per strain) measured from independent samples. Statistical significance was determined using a one-way ANOVA followed by Tukey’s multiple comparisons test (****, *P* < 0.0001).

**Figure S8.**
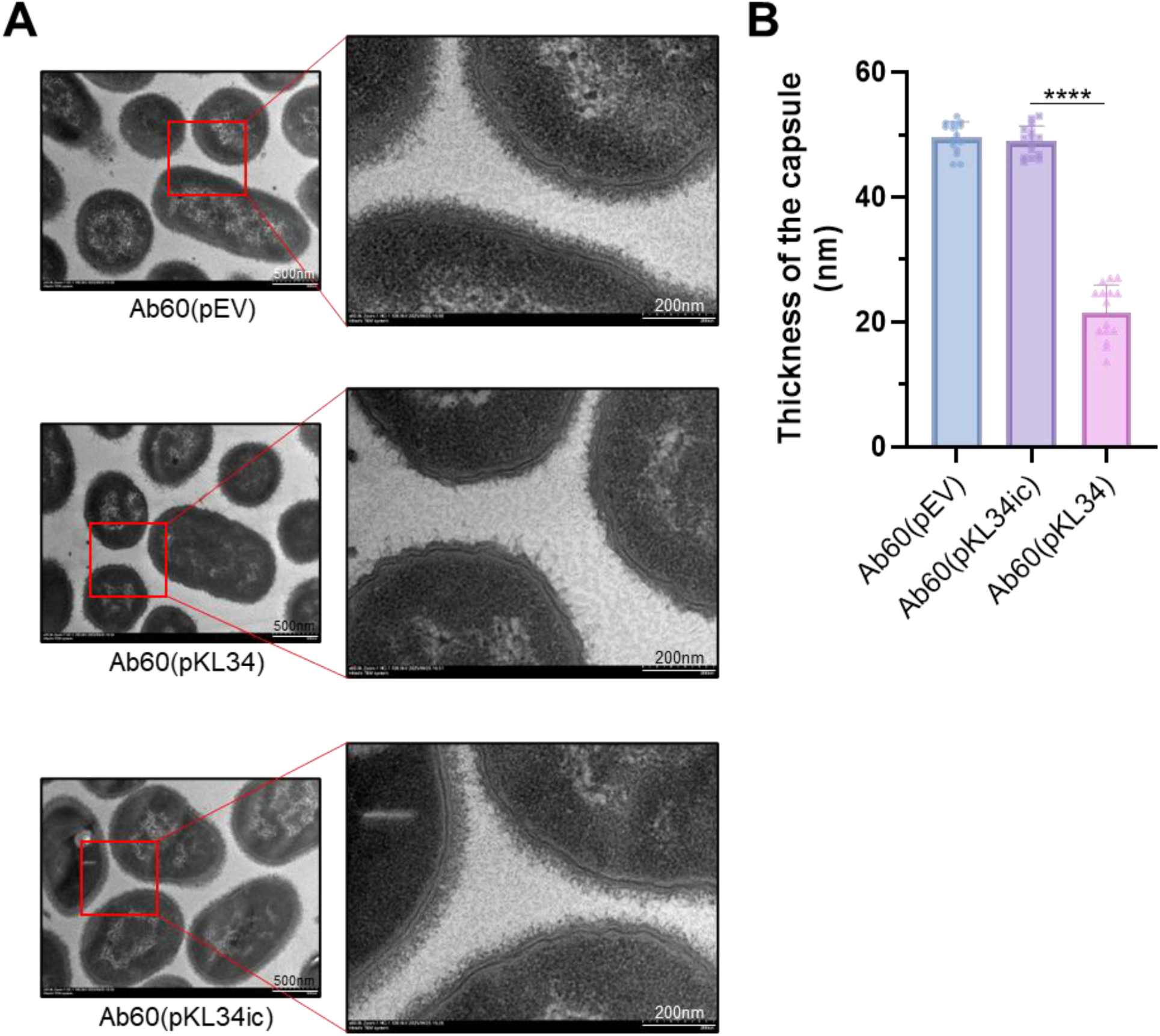
Heterologous expression of the KL34 gene cluster directs the formation of a high-density capsular shield. **(A)** TEM visualization of capsular morphology in Ab60-derived strains harboring the empty vector (pEV), pKL34ic, or pKL34. Only Ab60(pKL34) produced a dense capsular layer structurally resembling that of the naturally resistant strain Ab27. **(B)** Quantification of capsule thickness (upper panel) and relative electron density (lower panel). Data are presented as the mean ± SEM of individual bacterial cells (n ≥ 10 per condition). Statistical significance was determined using a one-way ANOVA followed by Tukey’s multiple comparisons test (****, *P* < 0.0001).

**Figure S9.**
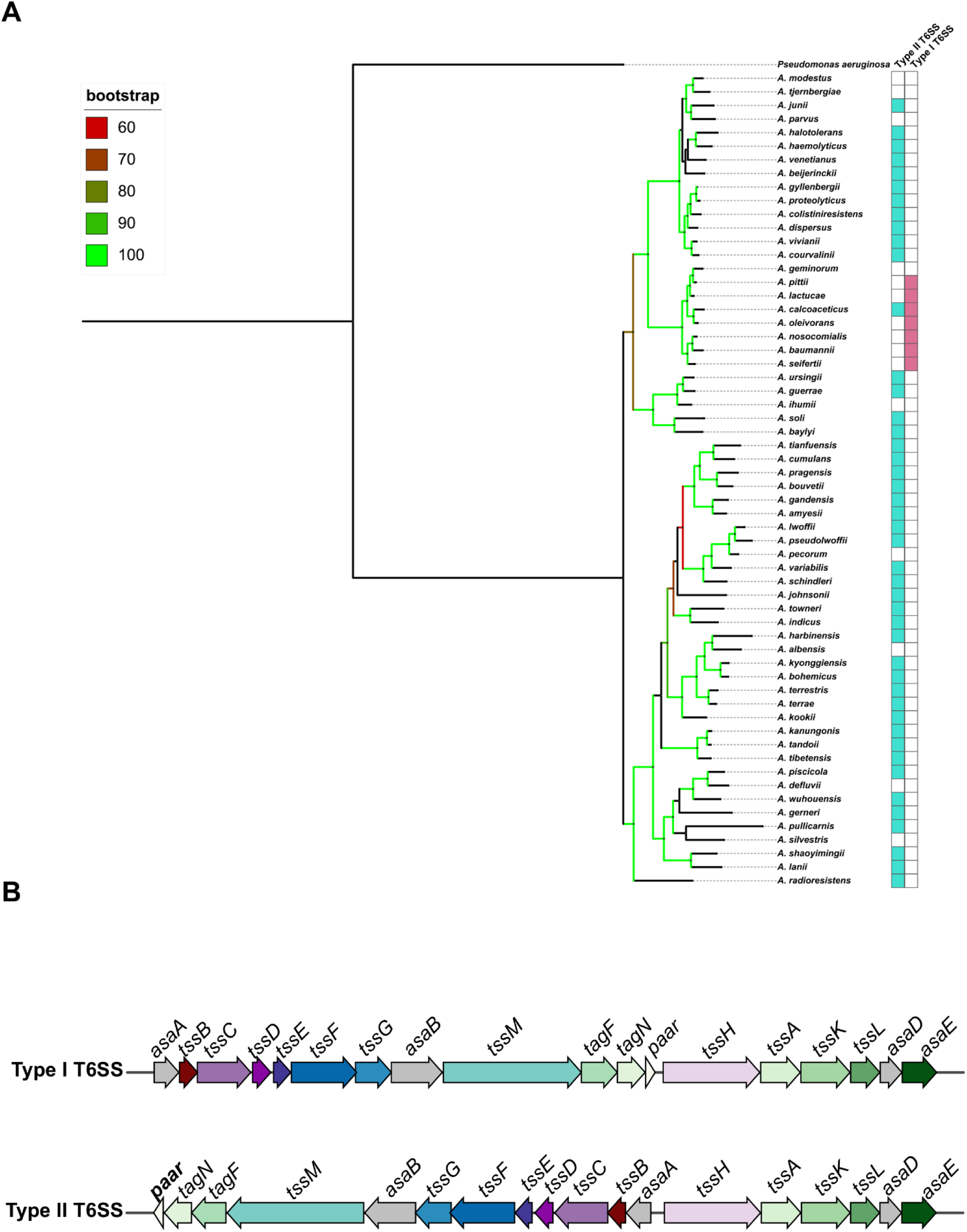
Phylogenetic divergence and distinct genetic organization of T6SS loci between pathogenic and nonpathogenic *Acinetobacter* species. **(A)** Phylogenetic relationships among pathogenic (e.g., *A. baumannii*, *A. nosocomialis*, *A. pittii*) and nonpathogenic *Acinetobacter* species based on core genome alignments. The adjacent heatmap indicates the distribution of two distinct T6SS structural subtypes: Type I and Type II. **(B)** Schematic comparison of the genetic architectures of T6SS subtypes. The Type I T6SS (top), conserved across pathogenic species, displays a uniform organization where all structural genes are transcribed unidirectionally. In contrast, the Type II T6SS (bottom), predominantly found in nonpathogenic species, exhibits a divergent arrangement where the first 12 genes upstream of *tssH* are transcribed in the opposite orientation.

**Figure S10.**
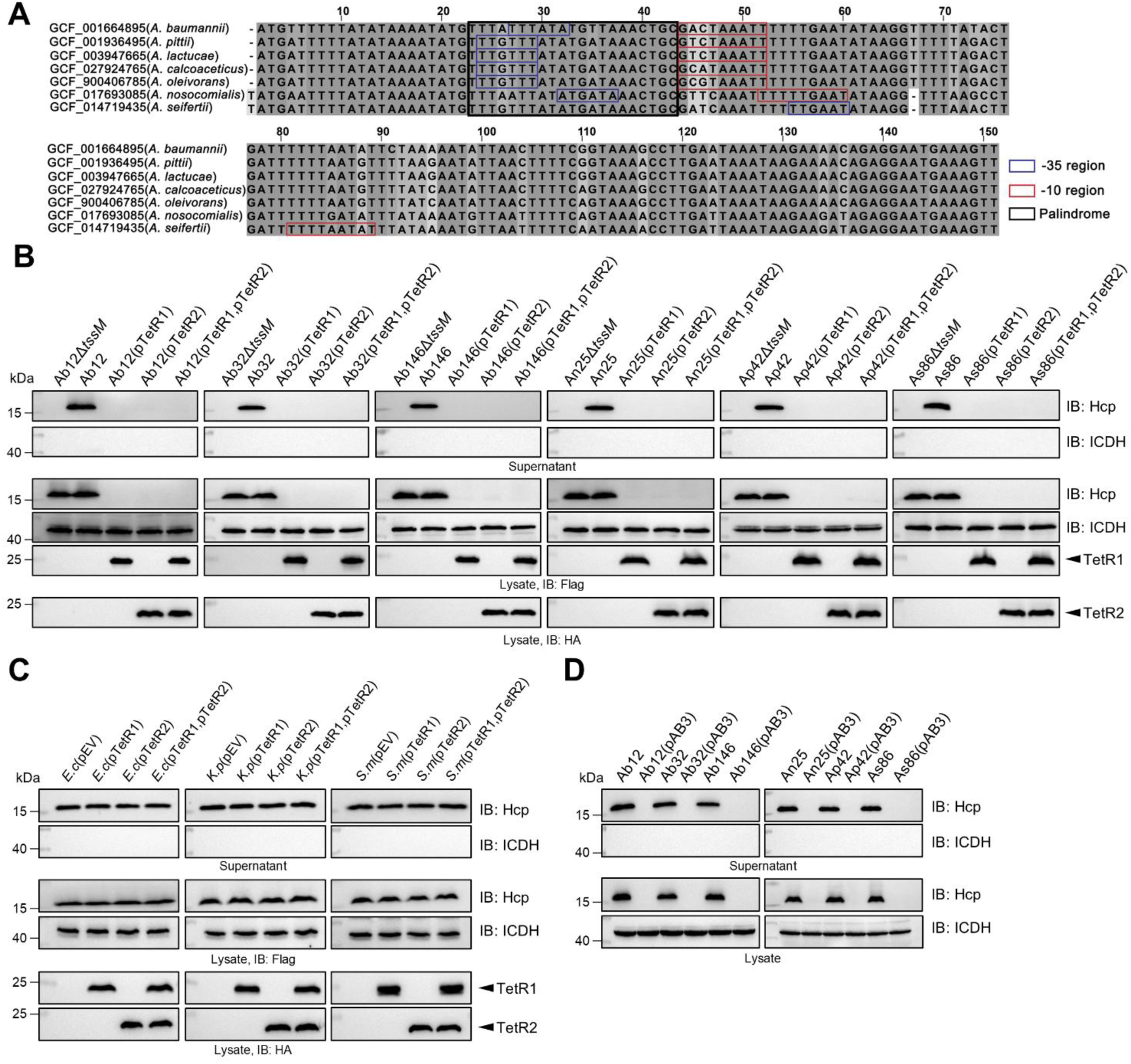
Conserved promoter architectures dictate the universal silencing of T6SS by pAB3-encoded TetR repressors across pathogenic *Acinetobacter*. **(A)** Multiple sequence alignment of the promoter regions governing Type I T6SS gene clusters in pathogenic *Acinetobacter* species. These regions share a highly conserved architecture, including specific –35 and –10 elements and a palindromic motif, indicating a universal regulatory mechanism. **(B)** Ectopic expression of TetR-family repressors broadly silences T6SS activity. Culture supernatants from diverse clinical isolates of *A. baumannii*, *A. nosocomialis*, *A. pittii*, and *A. seifertii* expressing TetR1 or TetR2 were probed for Hcp via immunoblotting. **(C)** The *Acinetobacter* plasmid-encoded repressors do not inhibit T6SS activity in distant Gram-negative competitors. Ectopic expression of TetR1 or TetR2 in *E. cloacae* and *K. pneumoniae* failed to suppress Hcp secretion. **(D)** Natural acquisition of the conjugative plasmid pAB3 abolishes host T6SS activity. Following the T4SS-mediated conjugative transfer of pAB3 into the indicated pathogenic recipient strains, Hcp secretion was assessed by immunoblotting.

**Figure S11.**
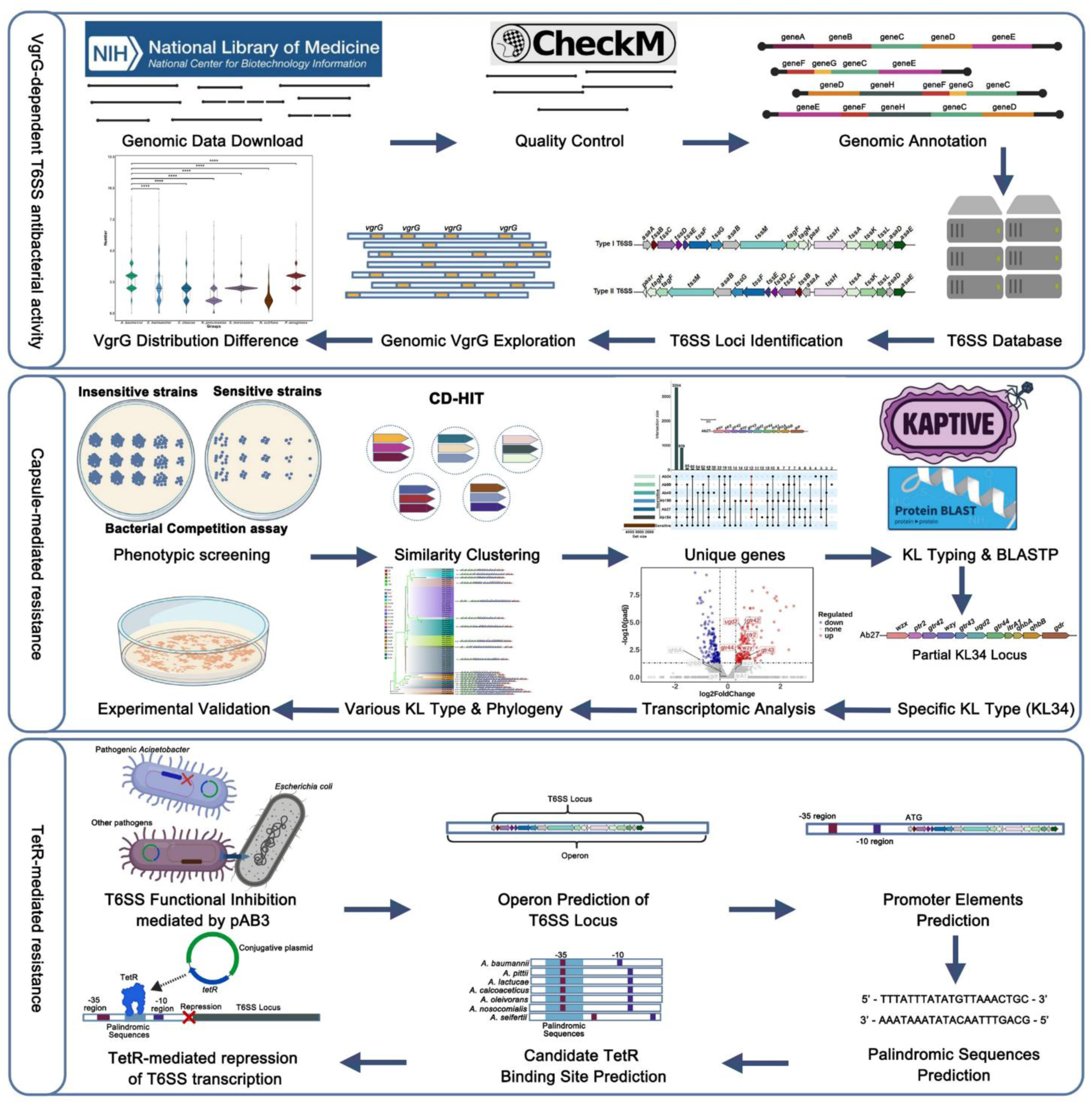
A schematic overview of the complete bioinformatic workflow and data processing pipeline.

